# Alternative ergosterol biosynthetic pathways confer antifungal drug resistance in the human pathogens within the *Mucor* species complex

**DOI:** 10.1101/2023.12.01.569667

**Authors:** María Isabel Navarro-Mendoza, Carlos Pérez-Arques, Josie Parker, Ziyan Xu, Steven Kelly, Joseph Heitman

## Abstract

Mucormycoses are emerging fungal infections caused by a variety of heterogeneous species within the Mucorales order. Among the *Mucor* species complex, *Mucor circinelloides* is the most frequently isolated pathogen in mucormycosis patients and despite its clinical significance, there is an absence of established genome manipulation techniques to conduct molecular pathogenesis studies. In this study, we generated a spontaneous uracil auxotrophic strain and developed a genetic transformation procedure to analyze molecular mechanisms conferring antifungal drug resistance. With this new model, phenotypic analyses of gene deletion mutants were conducted to define Erg3 and Erg6a as key biosynthetic enzymes in the *M. circinelloides* ergosterol pathway. Erg3 is a C-5 sterol desaturase involved in growth, sporulation, virulence, and azole susceptibility. In other fungal pathogens, *erg3* mutations confer azole resistance because Erg3 catalyzes the production of a toxic diol upon azole exposure. Surprisingly, *M. circinelloides* produces only trace amounts of this toxic diol and yet, it is still susceptible to posaconazole and isavuconazole due to alterations in membrane sterol composition. These alterations are severely aggravated by *erg3*Δ mutations, resulting in ergosterol depletion and consequently, hypersusceptibility to azoles. We also identified Erg6a as the main C-24 sterol methyltransferase, whose activity may be partially rescued by the paralogs Erg6b and Erg6c. Loss of Erg6a function diverts ergosterol synthesis to the production of cholesta-type sterols, resulting in resistance to amphotericin B. Our findings suggest that mutations or epimutations causing loss of Erg6 function may arise during human infections, resulting in antifungal drug resistance to first-line treatments against mucormycoses.

**Importance:** The *Mucor* species complex comprises a variety of opportunistic pathogens known to cause mucormycosis, a potentially lethal fungal infection with limited therapeutic options. The only effective first-line treatments against mucormycosis consist of liposomal formulations of amphotericin B and the triazoles posaconazole and isavuconazole, all of which target components within the ergosterol biosynthetic pathway. This study uncovered *M. circinelloides* Erg3 and Erg6a as key enzymes to produce ergosterol, a vital constituent of fungal membranes. Absence of any of those enzymes leads to decreased ergosterol and consequently, resistance to ergosterol-binding polyenes such as amphotericin B. Particularly, losing Erg6a function pose a higher threat as the ergosterol pathway is channeled into alternative sterols similar to cholesterol, which maintain membrane permeability. As a result, *erg6a* mutants survive within the host and disseminate the infection, indicating that Erg6a deficiency may arise during human infections and confer resistance to the most effective treatment against mucormycoses.

## Introduction

Mucorales comprise an order of human pathogenic filamentous fungi that includes *Mucor, Rhizopus*, and *Lichtheimia* species. These species cause infections known as mucormycoses, which typically occur when individuals inhale the spores of these molds or are infected through skin injuries, burns, or other traumatic wounds (1). Immunocompromised patients, such as those with cancer and/or undergoing immunosuppressive treatment, are particularly susceptible to invasive mucormycoses. Individuals suffering from poorly controlled diabetes mellitus are also at higher risk. This disease is associated with poor prognoses and dramatically high morbidity, resulting in mortality rates ranging from 23 % to 80 % (2). Because Mucorales are understudied pathogens, it is challenging to gather information on global incidence rates. Nevertheless, the trend over the past decade has shown an increase in the number of cases, especially associated with COVID-19 infections (3, 4), which has led the World Health Organization to include Mucoralean pathogens on their fungal priority pathogens list (5).

Recent global practice guidelines discourage the use of fluconazole, voriconazole (6), or the echinocandins (7) as therapeutic options for mucormycoses due to inherent resistance to those antifungal drugs. Instead, first-line treatment with liposomal formulations of the polyene amphotericin B is recommended at high doses, followed by salvage treatment with the triazoles posaconazole or isavuconazole (2). However, the applicability of these treatments can be limited by toxicity and resistance. Due to the limited availability of effective antifungal drugs, the emergence of resistant strains in the environment or in the host poses a global threat to public health. Antifungal drug resistance mechanisms in mucoralean clinical isolates are poorly understood, due to the lack of proper diagnostic tools and the heterogeneity of infections (5). Recent studies have revealed gain-of-function mutations in *erg11* confer resistance to azoles (6). Additionally, a novel mechanism for antifungal drug resistance based on RNA interference (RNAi), called epimutation, has been discovered in *Mucor* species (8). This mechanism could represent a source of transient antifungal drug resistance during host infection (9). The epimutation mechanism operates through RNAi-mediated gene silencing, repressing the expression of drug targets. Therefore, identifying loss-of-function mutations conferring antifungal drug resistance might elucidate new targets of this epigenetic process.

Understanding the mechanisms of action of the recommended antifungal therapies against mucormycoses may help to identify drug targets involved in drug resistance. Polyenes bind directly to ergosterol and both increase membrane permeability and sequester ergosterol (10, 11), while azoles block the main ergosterol biosynthetic pathway by inhibiting the Erg11 enzyme (12). By antagonistically binding Erg11, azoles can divert ergosterol biosynthesis into a different pathway that results in toxic sterol intermediates (13, 14). Thus, the mechanisms of action of both polyenes and azoles converge on ergosterol production and conceivably, altering the ergosterol biosynthetic pathway may lead to changes in drug susceptibility. Indeed, loss-of-function mutations in two key ergosterol biosynthetic genes, *erg3* and *erg6*, have been reported to result in antifungal drug resistance in other fungal pathogens. Erg3 is a C-5 sterol desaturase involved in the main ergosterol biosynthetic pathway, but also in the toxic sterol pathway (13). Therefore, loss of Erg3 function prevents the formation of toxic sterol intermediates produced during azole treatment, enabling evasion of the fungistatic effect of the drug (14, 15). On the other hand, Erg6 is a C-24 sterol methyltransferase that either converts zymosterol into fecosterol or lanosterol into eburicol (16), two alternative ergosterol precursors. Mutations affecting Erg6 function favor the production of cholesta-type sterols, conferring polyene resistance (17–23).

Research on molecular mechanisms leading to drug resistance in Mucorales is of paramount importance, albeit challenging given the current genetic and molecular models and tools. *Mucor* is a genus of related fungal species within the order Mucorales. *M. circinelloides* is the *Mucor* species most frequently isolated from clinical sources and is associated with mucormycotic infections (24–26). Previously, the *Mucor circinelloides* species complex was delineated into *formae* but recent taxonomic and phylogenetic analyses have defined a minimum of 16 distinct phylogenetic species, 7 of which are known to cause human disease (27). As a consequence, research has advanced genetics, genomics, and hostmicrobe interaction models for this understudied group of pathogens, leading to the development of *Mucor* as a model for mucormycoses over the past decade (8, 28–31). Most of these studies relied on *M. lusitanicus* because it was the only species amenable to genetic transformation and it has a well assembled genome (32, 33). Yet among *Mucor* species, *M. lusitanicus* is not a frequent cause of mucormycosis, probably due to its maximum growth temperature being 35 to 37 °C, compared to *M. circinelloides*, which grows at 37 to 39 °C (27). In this study, we introduce a new model to study mucormycosis pathogenesis and antifungal drug susceptibility in *M. circinelloides*. We developed a novel transformation method to characterize the *erg3* and *erg6* genes in a *M. circinelloides* pathogenic isolate, leading to the discovery of new mechanisms for amphotericin B resistance and azole susceptibility in the Mucorales.

## Results

### A genetic transformation procedure for *Mucor circinelloides* pathogenic species

*M. circinelloides* is the most frequent *Mucor* species isolated from mucormycosis patients, and these isolates are usually thermotolerant and highly virulent in host-pathogen infection models, yet reliable genetic transformation procedures are not available for this species. To develop a genetic model for virulence and antifungal susceptibility studies, we aimed at generating a *M. circinelloides* strain that is competent for genetic transformation. Because ensuring full virulence and thermotolerance in our model was a priority, *M. circinelloides* 1006PhL human isolate (hereinafter *M. circinelloides*) was selected to conduct studies as it has both attributes (34). Also, its genome has been sequenced (35) and is publicly available (https://www.ncbi.nlm.nih.gov/datasets/genome/GCA_000401635.1/), providing an advantage to genetic manipulation. Traditionally in mucoralean genetics, the application of dominant resistance markers is not possible due to high intrinsic resistance to these drugs. *M. circinelloides* is not an exception, as it exhibits residual growth upon exposure to drugs frequently utilized for transformant selection, complicating the discrimination between resistant transformants and residually growing colonies (Figure S1).

As a consequence, we designed a workflow to obtain an auxotrophic mutant that could serve as a transformation recipient strain, based on successful previous studies (36). 5-fluoroorotic acid (5-FOA) is sequentially converted into cytotoxic 5-fluorouracil (5-FU) by the action of the orotate phosphoribosyltransferase PyrF and orotidine-5’-monophosphate decarboxylase PyrG. As Mucorales are haploid fungi, and these two enzymes catalyze essential steps in the uracil biosynthetic pathway, a single loss-of-function mutation in either of their encoding genes results in resistance to 5-FOA and uracil auxotrophy that can be rescued by uridine and/or uracil supplementation. Following this rationale (Figure 1A), we selected for spontaneous uracil auxotrophic mutants by utilizing 5-FOA but avoiding additional mutagenic agents to prevent undesired secondary mutations. A total of 237 5-FOA resistant colonies were isolated and 8 were tested for uracil auxotrophy by comparing their ability to grow on media with or without uridine and uracil. Among these, one isolate named MIN6 was unable to grow without uridine and uracil indicative of uracil auxotrophy (Figure 1B, isolate 6).

**Figure 1.**
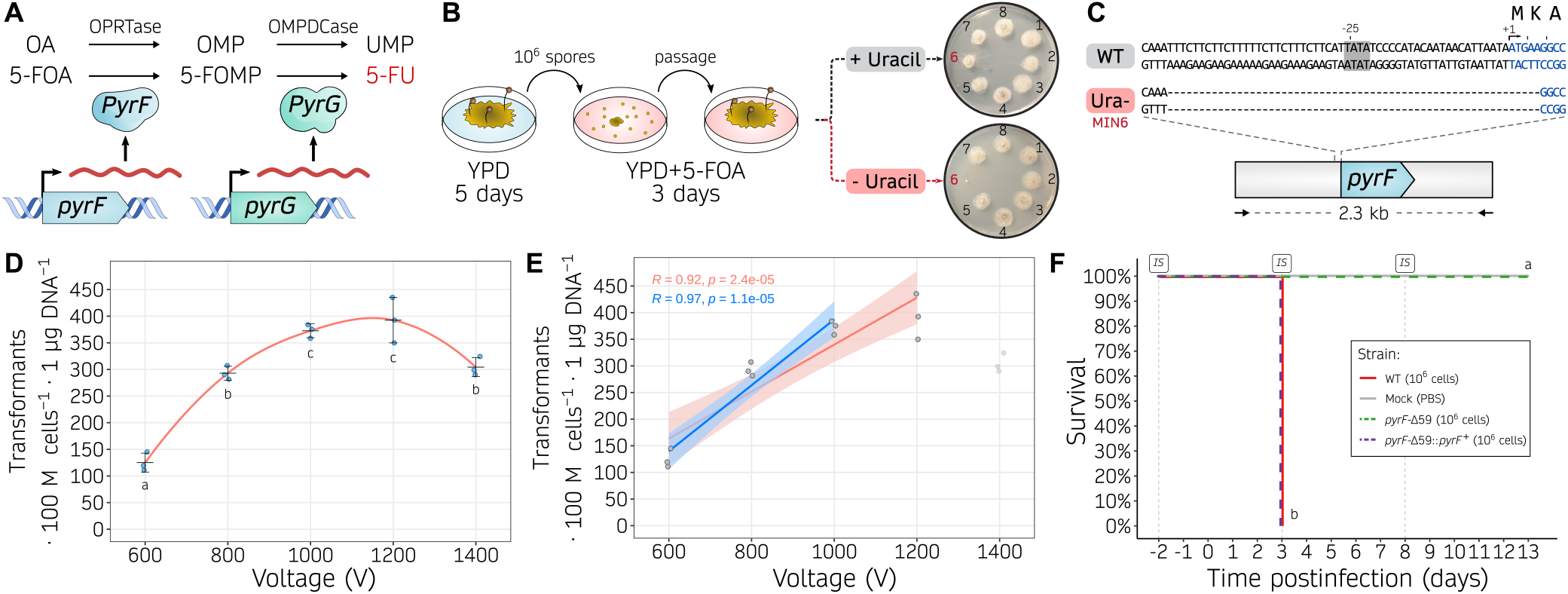
Uracil auxotrophy enables *M. circinelloides* transformation based on a *pyrF* selectable marker. **(A)** Role of PyrF (orotate phosphoribosyltransferase, OPRTase) and PyrG (orotidine monophosphate decarboxylase, OMPDCase) in the uracil biosynthetic pathway, converting orotic acid (OA) to orotidine monophosphate (OMP), and OMP into uridine monophosphate (UMP), respectively. Alternatively, the same enzymes catalyze the conversion of 5-fluoroorotic acid (5-FOA) into fluoro-OMP (FOMP), and FOMP into 5-fluorouracil (5-FU), respectively. **(B)** Experimental design followed to isolate uracil auxotrophic strains. Briefly, wildtype spores were collected from YPD medium, counted, and one million spores were plated onto YPD containing 5-FOA. After three days, 5-FOA resistant colonies were isolated and transferred to new 5-FOA containing medium to confirm the resistance, and then inoculated onto minimal medium either with or without uracil to confirm the auxotrophy. **(C)** *pyrF* sequences from the wildtype (WT) and uracil auxotrophic (Ura-, MIN6) strains, focusing on the 5’-region of the gene. The TATA box motif (gray background around position -25), transcription start site (arrow, position +1), *pyrF* coding sequence (blue font), and encoded one-letter amino acids are shown. **(D, E)** Transformation efficiency measured as transformants per 100 million protoplasts per microgram of DNA using different voltages in the electroporation procedure. Individual values (dots) from biological triplicates were used to determine the mean and SD values as black lines. In **(D)**, a smoothed curve is drawn as a red line. Significant differences among voltages were determined by a One-way ANOVA and Tukey HSD test, and voltages showing different letters indicate significant differences (*p-value* ≤ 0.05). In **(E)**, two linear regression models are displayed showing a positive correlation for voltages from 600 to 1,000 V (blue line) or to 1,200 V (red line). Pearson’s correlation coefficients (R) are shown for each regression. **(F)** Kaplan-Meier survival curves of immunosuppressed mice after infection with *M. circinelloides* wildtype and transformable strains (color-coded). Intraperitoneal immunosuppressive (IS) treatments are shown. Strains showing significant differences in virulence are indicated as different letters (*p-value* ≤ 0.0001), assessed by a Log-rank test.

Subsequent Sanger-sequencing of the *pyrF* and *pyrG* genes revealed a 59-basepair (bp) microdeletion within the *pyrF* locus (*pyrF*-Δ59). This microdeletion comprises the last 54 bp of the promoter region, including a putative TATA-box element whose deletion may impair transcription; and the first 5 bp of the coding sequence, leading to a start loss and a consequent frameshift mutation assuming translation would start at the next available AUG triplet (Figure 1C).

Next, we complemented MIN6 uracil auxotrophy by integrating the *pyrF* marker, a 2.3-kilobase (kb) DNA fragment comprising the wildtype *pyrF* promoter, coding sequence, and terminator regions aiming at favoring homologous recombination and subsequent replacement of the *pyrF*-Δ59 allele with its wildtype sequence. To integrate the *pyrF* marker, we improved a reliable protoplast transformation procedure involving electroporation. For optimal protoplast preparation, the spore germination rate in *M. circinelloides* was comparatively slower than in *M. lusitanicus*; this delay may be attributed to the pre-swollen and larger size of *M. lusitanicus* spores (37). DNA was introduced into the protoplasts by exponential decay waveform electroporation with constant capacitance and resistance, and transformation efficiency was assessed at different voltages (Figure 1D). The most efficient transformation rates were obtained with 1000 and 1200 V. A linear regression modeling indicates a positive correlation between transformation efficiency and voltage, particularly significant at voltages ranging from 600 to 1000 V (Figure 1E). Therefore, subsequent electroporations were carried out at 1000 V to ensure maximum efficiency while avoiding unnecessary protoplast lethality. The successful complementation of the *pyrF*-Δ59 allele was verified by PCR-amplification and restriction fragment length polymorphism (RFLP, Figure S2).

Next, we determined the effect of the *pyrF*-Δ59 allele on virulence and tested whether *pyrF* complementation could rescue a fully virulent phenotype (Figure 1F). To do so, the wildtype 1006PhL strain, the *pyrF*-Δ59 uracil auxotrophic strain, and the *pyrF*-Δ59::*pyrF*^+^ complemented strain were analyzed in a survival assay using immunosuppressed mice. The uracil auxotrophic strain showed a complete and significant lack of virulence, whereas the wildtype 1006PhL and *pyrF*^+^ complemented strain displayed full virulence without significant differences. Similarly, fungal virulence correlated with animal weight loss (Figure S3). These results demonstrate that uracil auxotrophy impairs the pathogenic potential of *M. circinicelloides*, which can be restored by *pyrF*^+^ complementation. Overall, these findings highlight this procedure as a highly efficient, reliable method to obtain uracil auxotrophic strains, transform by electroporation, and replace genes by homologous recombination in *M. circinelloides*. But more importantly, they support the application of this genetic model to study virulence processes and antimicrobial drug resistance within the *Mucor* species complex, as opposed to other thermosensitive, partially virulent, and clinically less relevant species such as *M. lusitanicus*.

### Erg3 and Erg6 distribution across the Mucoromycota

We aimed to delete the *erg3* and *erg6* genes in *M. circinelloides* and *M. lusitanicus* to illustrate the potential of our newly transformable *M. circinelloides* strain for standardized antifungal susceptibility testing and virulence assays. A protein-protein similarity search identified one Erg3 and three Erg6 homologs whose evolutionary history was examined in several pathogenic species from the Mucoromycota, as well as in model organisms with single-copy *erg* genes for reference (Figure 2, Table S1 for Erg3, and Table S2 for Erg6 sequences). Erg3 protein sequences from Mucoromycota fungi form a robust clade within fungal Erg3 proteins, highly consistent with their species phylogeny (Figure 2A). This phylogeny suggests two independent duplication events in the Lichtheimiaceae and Rhizopodaceae families as most of their members harbor two Erg3 paralogs, which could be explained by recent segmental duplication events in these families (38–40). Besides these specific duplication events, most Mucoromycota species harbor a single Erg3 ortholog and particularly, the *Mucor* clade. The Erg3 ortholog exhibits high synteny among *Mucor* species (Figure 2C and Figure S4A), confirming our phylogenetic inference. In addition, transcriptomic analysis revealed that *erg3* is actively transcribed in both *Mucor* species. Taken together, these results indicate that *erg3* deletion will suffice to confer loss of Erg3 function.

**Figure 2.**
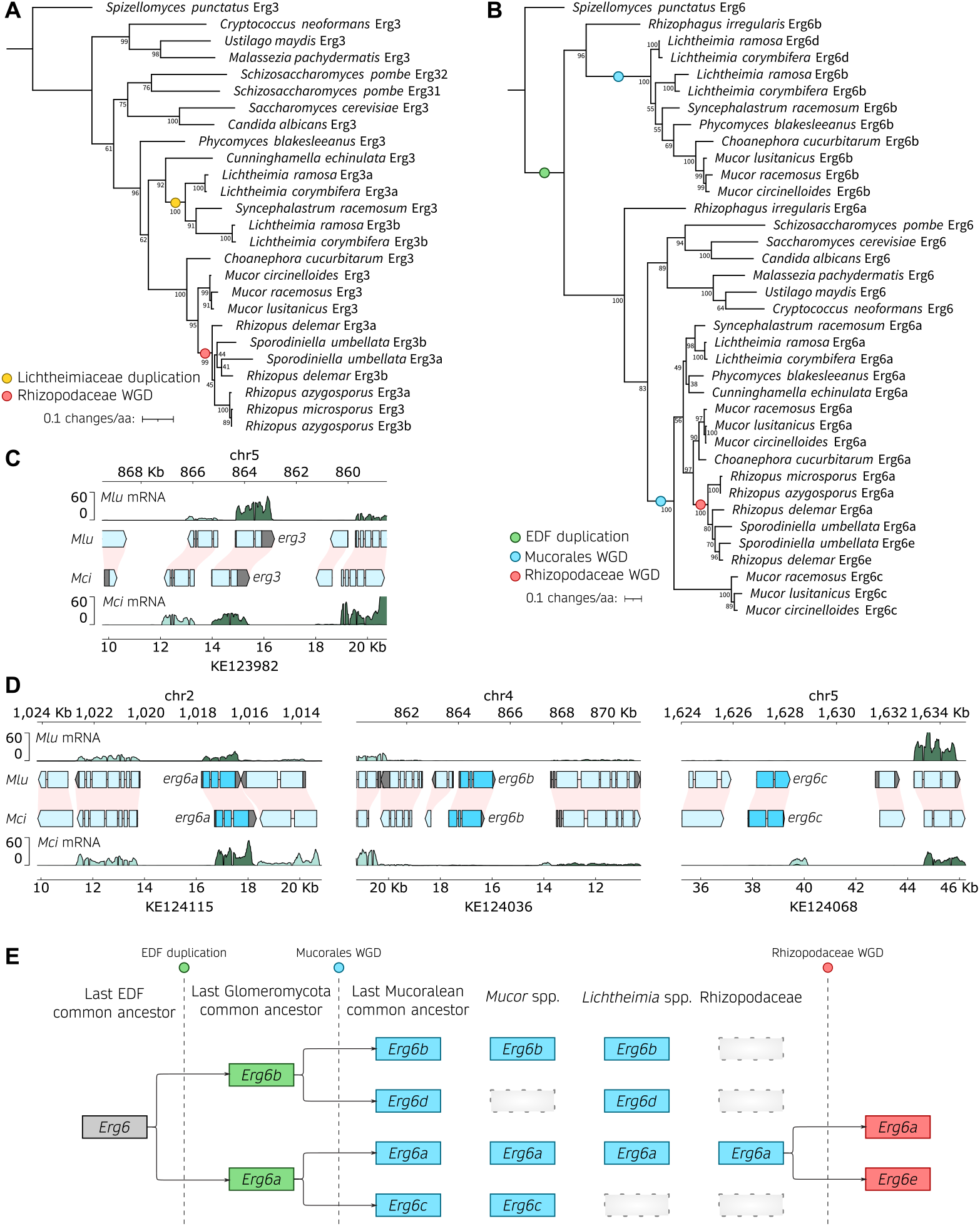
Erg3 and Erg6 evolutionary history across the Mucoromycota. **(A, B)**. Phylogenetic tree depicting the Erg3 **(A)** and Erg6 **(B)** amino acid sequence alignments in clinically relevant fungi, with a specific focus on Mucorales. The Erg protein sequences from the chytrid *Spizellomyces punctatus* were employed as outgroups to root the trees. Additionally, the Erg6 sequence from *Rhizophagus irregularis*, a member of the Glomeromycota and the closest non-Mucoromycota sequence, was included to provide clarity on duplication events in **(B)**. Noteworthy branching events, including whole genome duplications (WGD) or gene duplications, are denoted by color-coded circles. Node support is represented as a percentage based on 1,000 bootstrapping iterations. The branch length is scaled to 0.1 substitutions per amino acid, as indicated in the legend. **(C, D)**. Plots illustrating gene transcription across *M. lusitanicus* (*Mlu*) and *M. circinelloides* (*Mci*) genomic regions containing *erg3* **(C)** and *erg6* orthologs **(D)**. The genomic coordinates for both species are displayed above (*Mlu*) and below (*Mci*) each plot. Gene annotation is depicted as arrowed blocks, indicating the direction of transcription and the boundaries of exons and introns. The coloring scheme assigns cyan blue to *erg* coding sequences, light blue to the remaining coding sequences, and gray to untranslated regions. Interspecies synteny among genes is depicted as pink shading. Gene transcription is shown as rRNA-depleted RNA read coverage, with the forward strand represented in dark green and the reverse strand in light green, enabling assessment of sense and antisense transcription. In some instances, the x-axes were inverted, and in those cases, the forward and reverse strand orientations were also inverted to enhance visualization and clarification. **(E)** Model illustrating the inferred evolutionary history of Erg6 across the early-diverging fungi (EDF).

On the other hand, the evolutionary history of Erg6 is more complex (Figure 2B). We identified a robustly supported, monophyletic clade encompassing the orthologs from well-known model organisms as well as Glomeromycota and Mucoromycota Erg6a paralogs. This clade exhibits two different Erg6 paralogs from *Mucor* species: one closer to the clade root and the reference Erg6 sequences (Figure 2B, Erg6a) and another considerably divergent (Figure 2B, Erg6c, bottom clade). In addition, most species from the Mucoromycota harbor another Erg6 paralog, that is more divergent compared to the reference Erg6 sequences (Figure 2B, Erg6b). We developed a model (Figure 2E) that offers the most plausible interpretation of the phylogenetic and synteny patterns observed across all of the Mucoromycota species analyzed, explaining the origin of these paralogs (Figure S4D-H). An ancient duplication event predating the Glomeromycota and Mucoromycota common ancestor resulted in two highly divergent Erg6 paralogs in species from the Mucoromycota and the glomeromycete *Rhizophagus irregularis* (Figure 2B, EDF duplication, Erg6a and Erg6b). Subsequent genome duplications in the mucoralean common ancestor (32), along with gene loss, led to most Mucoromycota species harboring two to three Erg6 paralogs. Interestingly, gene loss did not occur uniformly in all clades: e.g., the third paralog found in *Mucor* species (Erg6c) is different from that of *Lichtheimia* species (Erg6d), and several species harbor only two paralogs (Erg6a and Erg6b). Moreover, Rhizopodiaceae species exhibit an additional, more recent duplication (Erg6e), which is consistent with another round of whole-genome duplication and subsequent gene loss in this lineage (39, 40). All three Erg6 paralogs found in *Mucor* species, which also show conserved synteny (Figure 2D and Figure S4D-H), are explained by this model. Interestingly, only *erg6a* is actively expressed (Figure 2D), which together with the phylogenetic data suggests that *erg6b* and *erg6c* may not be as biologically relevant during standard growth conditions. Therefore, we named the *Mucor* paralogs Erg6a, Erg6b, and Erg6c in agreement with our evolutionary model (Figure 2E), phylogenetic, and transcriptomic data, and we reasoned that *erg6a* deletion may suffice to confer loss of Erg6 function.

To explore the role of the *erg3* and *erg6a* genes in mucoralean ergosterol biosynthesis, antifungal susceptibility, and pathogenesis, we generated deletion mutant strains in *M. circinelloides* and *M. lusitanicus* (Figure S5). The genetic loci were successfully replaced through homologous recombination with uracil prototrophy markers: *pyrF* for *M. circinelloides* (Figure S5A-B), utilizing the newly designed MIN6 strain and genetic transformation via electroporation procedure; and *pyrG* for *M. lusitanicus* (Figure S5C-D), using a previously described transformation method (41). Because Mucorales have non-septate hyphae and multinucleated spores, transformed protoplasts frequently harbor wild-type and mutant nuclei simultaneously, a phenomenon known as heterokaryosis. To lose the wildtype nuclei and obtain exclusively homokaryotic deletion mutants, transformants must be cultured for several asexual generations in a medium that positively selects the mutant nuclei. After this process of mutant nuclei enrichment, *erg3*Δ and *erg6a*Δ homokaryotic mutations were verified by PCR-amplified fragment length polymorphism (AFLP, Figure S5), and two independently generated homokaryotic mutants were selected for each deletion to ensure phenotypic specificity in subsequent studies.

### *erg3* and *erg6a* deletion affects growth and virulence in *M. circinelloides*

Alterations in ergosterol composition may affect the stability of biological membranes and usually lead to fitness impairments. Consequently, we examined the effects of loss-of-function mutations in the *erg3* and *erg6a* genes of *M. circinelloides* and *M. lusitanicus*, specifically focusing on evaluating defects in various aspects of fitness, including growth rate, sporulation, and virulence. For these and subsequent analyses, data from each pair of independently generated mutants harboring the same gene deletion were grouped for simplicity when necessary for primary figures, after ensuring that there were not significant differences between them as can be observed in the supplementary information (Figures S6-S8, and Data set S1). The growth rate was evaluated at different temperatures, including optimal growth and human physiological temperatures (Figure 3A,B). The *M. circinelloides erg3*Δ mutant exhibited a significant growth delay under all conditions tested (26 C, 30 ^*°*^C, and 37 ^°^C) compared to the wild type, whereas the *M. circinelloides erg6a*Δ mutation did not affect growth significantly (Figure 3A). The same growth trends were observed in minimal media (Figure S6A). Similarly, the *M. lusitanicus erg3*Δ mutant displayed growth defects under every condition (26 ^°^C and 30 ^°^C), while the *erg6a*Δ mutant showed a growth delay only at 30 ^°^C (Figure 3B). As *M. lusitanicus* is not able to withstand human physiological temperatures (37 ^°^C), and exhibits a considerably lower optimal growth temperature (26 ^°^C), this could explain the interspecies differences observed. Notably, increasing the temperature range beyond the optimal growth temperature for each species (37 ^°^C for *M. circinelloides* and 30 ^°^C for *M. lusitanicus*) severely affected the growth rate in every strain and wildtype controls but more importantly, rendered the phenotype confirmed by *erg* gene deletions more severe. All mutant strains exhibited full uracil prototrophy after *pyrF* or *pyrG* insertion, and the leucine auxotrophy in *M. lusitanicus* did not affect their growth rates, as no differences were observed upon uridine, uracil, and leucine supplementation (Figure S6C,E).

**Figure 3.**
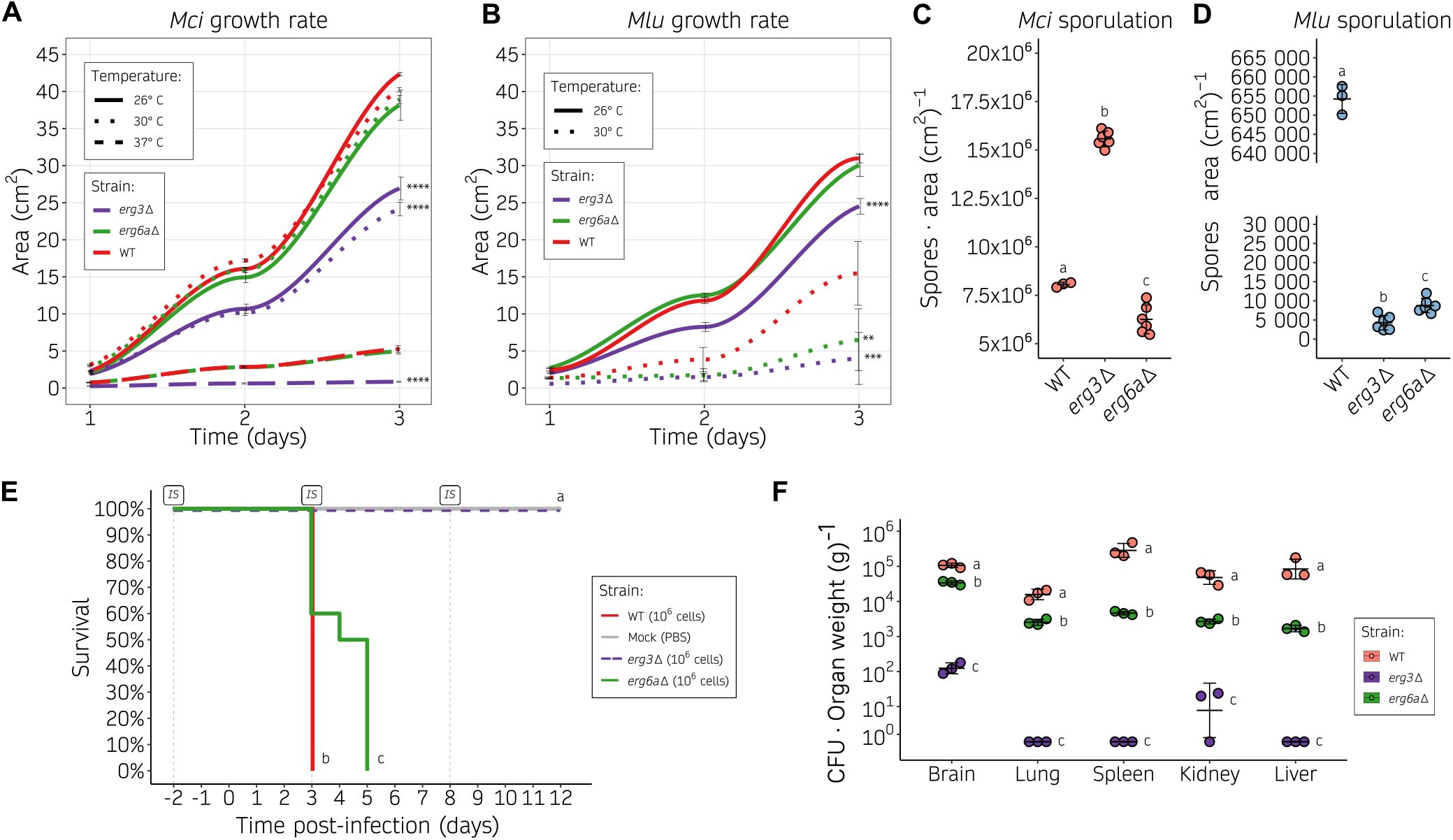
Alterations in the ergosterol biosynthetic pathway result in decreased growth rate, sporulation, and virulence. **(A, B)** Growth rate of *M. circinelloides* **(A)** and *M. lusitanicus* **(B)** *erg3*Δ and *erg6a*Δ (shown as color-coded lines) on YPD medium at different temperatures (shown as distinct line types), measured as colony area across 24-hour intervals. Individual values were determined either in six biological replicates for the wild type or in three biological replicates of two independently generated mutants (six total values, grouped together for simplicity as no significant differences were detected), and used to plot a smoothed curve and SD values as black lines. Significant area differences in depicted strains compared to the wildtype growth rate across the whole time-course were determined by a One-way ANOVA and Tukey HSD test for each temperature group and shown as asterisks (^**^ *p-value* ≤ 0.01, ^***^ *p-value* ≤ 0.001, and ^****^ *p-value* ≤ 0.0001). **(C, D)**. Sporulation rate of *M. circinelloides* **(C)** and *M. lusitanicus* **(D)** *erg3*Δ and *erg6a*Δ after a 72-hour incubation at room temperature on YPG medium, determined as number of spores per cm^2^. Values from individual measurements were obtained from three biological replicates of either the wildtype strain or two independently generated mutants (grouped together as no significant differences were found). These values are depicted in a dot plot, with standard deviation (SD) values represented as black lines. Strains are grouped by letters showing significant sporulation differences determined by a One-way ANOVA and Tukey HSD test (*p-value* ≤ 0.01). **(E)** Kaplan-Meier survival curves of immunosuppressed mice after infection with *M. circinelloides* wildtype and ergosterol mutants (color-coded). For *erg3*Δ and *erg6a*Δ, two independently generated mutants were injected into groups of 5 mice. For simplicity, after observing no significant differences in the results, survival curves were grouped together per gene deletion in groups of 10. Intraperitoneal immunosuppressive (IS) treatments are shown. Strains showing significant differences in virulence are indicated as different letters (*p-value* ≤ 0.01), assessed by a Log-rank test. **(F)** Fungal burden at 3 days post-infection in immunosuppressed mice, measured as colony-forming units (CFU) per gram of five different organs. SD and mean (black lines) were assessed from 3 biological triplicates. For each organ category, strains are grouped by letters showing significant fungal burden differences (One-way ANOVA and Tukey HSD test, *p-value* ≤ 0.01).

Asexual sporulation was also affected by *erg* mutations. In *M. circinelloides*, the *erg3*Δ mutant displayed a significant increase in sporulation, as opposed to the decrease observed in the *erg6a*Δ mutant compared to the wildtype strain (Figure 3C). On the other hand, *M. lusitanicus erg3*Δ and *erg6a*Δ mutants showed a severe and significant reduction in sporulation.

The observed differences in both growth and sporulation imply that deletion of the *erg3* and *erg6a* genes could impact fitness under various stress conditions, particularly temperature. To investigate whether these defects are relevant during the infection process, a survival assay was conducted using *M. circinelloides* strains, as they withstand murine physiological temperatures and exhibit a more reliable pathogenic potential. Immunosuppressed mice were infected with two independently generated mutants for each gene, and with the wildtype control strain (Figure 3E and Figure S7). Although *erg6a*Δ mutants exhibited attenuated virulence compared to the wildtype control, they were able to establish a fully disseminated infection in every organ analyzed for fungal burden, resulting in mortality among all infected mice. Conversely, animals infected with *erg3*Δ mutants survived, possibly due to the significant and drastic reduction in fungal burden within vital organs (Figure 3E-F). These findings suggest that *erg3* deletion abolishes virulence in *M. circinelloides*, whereas *erg6a* deletion may not suffice to prevent dissemination and mortality.

Taken together, these results confirm that deletion of *erg3* and *erg6a* leads to a reduction in fitness in specific settings and particularly at higher than optimal temperatures, observing a higher fitness cost for *M. lusitanicus*, and confirming interspecies differences between *M. circinelloides* and *M. lusitanicus*. As expected, this *erg* deletion-related loss of fitness impacts the virulence potential of *M. circinelloides*, more profoundly as a consequence of *erg3* deletion, and slightly due to *erg6a* deletion. Although *erg6a* deletion leads to delayed infection, it is noteworthy that this strain is still able to cause full mortality.

### *erg6a* loss-of-function mutations result in antifungal drug resistance

The antifungal drug susceptibility profiles of *erg3*Δ and *erg6a*Δ mutants were conducted following the CLSI and EUCAST standard methodology for filamentous fungi, determining the minimal inhibitory concentration values required to inhibit growth (MIC) by broth microdilution (Table 1). Susceptibility testing focused on the only three clinical antifungal drugs recommended for mucormycosis treatment: the liposomal formulation of amphotericin B (Ambisome), posaconazole (Noxafil), and isavuconazonium sulfate (Cresemba). Currently, clinical breakpoints have not been established for any of these treatments for Mucorales and therefore, it is not possible to strictly assess resistance or susceptibility. However, for clarity, the term “resistance” will henceforth refer to decreased susceptibility in our isolates. Both *M. circinelloides* and *M. lusitanicus* exhibited similar trends in MIC values, indicating generalizable alterations in antifungal drug susceptibilities. First, *erg3* deletion caused a minor increase in MIC for amphotericin B, with a 2-fold and 4-fold increase observed in *M. circinelloides* and *M. lusitanicus*, respectively. Conversely, *erg3*Δ mutants also exhibited an increased susceptibility to azoles, as evidenced by a 4-fold decrease in MIC for posaconazole, and a 2-fold decrease for isavuconazole in both *M. circinelloides* and *M. lusitanicus*. Second, the e*rg6a*Δ mutants showed substantial resistance to amphotericin B, specifically a 4-fold and 16-fold MIC increase in *M. circinelloides* and *M. lusitanicus*, respectively. These *erg6a*Δ mutants also exhibited a slightly increased susceptibility to azoles, manifested by a 2-fold MIC decrease for posaconazole and isavuconazole in *M. lusitanicus*, and a 2-fold decrease for posaconazole in *M. circinelloides*.

**Table 1.** Amphotericin B (AMB), posaconazole (POS) and isavuconazole (ISA) minimal inhibitory concentration (MIC) for *M. circinelloides* and *M. lusitanicus erg3*Δ and *erg6a*Δ mutants.

| Strain | AMB (mg/L) | POS (mg/L) | ISA (mg/L) |
| --- | --- | --- | --- |
| *M. circinelloides* |  |  |  |
| WT 1006PhL | 0.125 | 2 | 8 |
| *erg3*Δ-1 | 0.25 | 0.5 | 4 |
| *erg3*Δ-2 | 0.25 | 0.5 | 4 |
| *erg6a*Δ-1 | 0.5 | 1 | 8 |
| *erg6a*Δ-2 | 0.5 | 1 | 8 |
| *M. lusitanicus* |  |  |  |
| WT MU636 | 0.03125 | 0.5 | 4 |
| *erg3*Δ-1 | 0.125 | 0.125 | 2 |
| *erg3*Δ-2 | 0.125 | 0.125 | 2 |
| *erg6a*Δ-1 | 0.5 | 0.25 | 2 |
| *erg6a*Δ-2 | 0.5 | 0.25 | 2 |

Antifungal drug susceptibility testing in molds by broth microdilution is challenging and subject to qualitative interpretation. More importantly, it is difficult to account for the inherent growth defects that are present in *erg3* and *erg6a* deletion strains. An alternative method was employed to address this issue, measuring antifungal drug susceptibility as the area of inhibited growth on solid media, normalizing to growth without drug. To obtain a dynamic quantification of drug susceptibility, area measurements were taken at different time points (Figure 4 and Figure S8). The differences in drug susceptibility became more evident on day 2 (Figure 4A), as indicated by a plateau in the growth inhibition curves between days 2 and 3 (Figure 4B-C). Growth inhibition was stronger on day 1 in every condition tested and stabilized after day 2, suggesting that amphotericin B and azoles also restrict germination in addition to growth. The *erg3*Δ mutants of *M. circinelloides* and *M. lusitanicus* showed an increased susceptibility to both azoles –posaconazole and isavuconazole– and this hypersusceptibility was more severe under isavuconazole stress. Similarly, amphotericin B resistance was confirmed in *erg6a*Δ mutants for both *Mucor* species, and in *M. circinelloides erg3*Δ mutants. *M. lusitanicus erg6a*Δ mutants also showed an increased susceptibility to azole drugs, overall confirming that these differences in susceptibility observed in the MIC tests (Table 1) were significant and independent of their inherent growth defects. Taken together, our results indicate that modifying the ergosterol biosynthetic pathway, and consequently, the content of ergosterol and intermediate sterols, may contribute to the observed drug susceptibility changes that we aim to address further.

**Figure 4.**
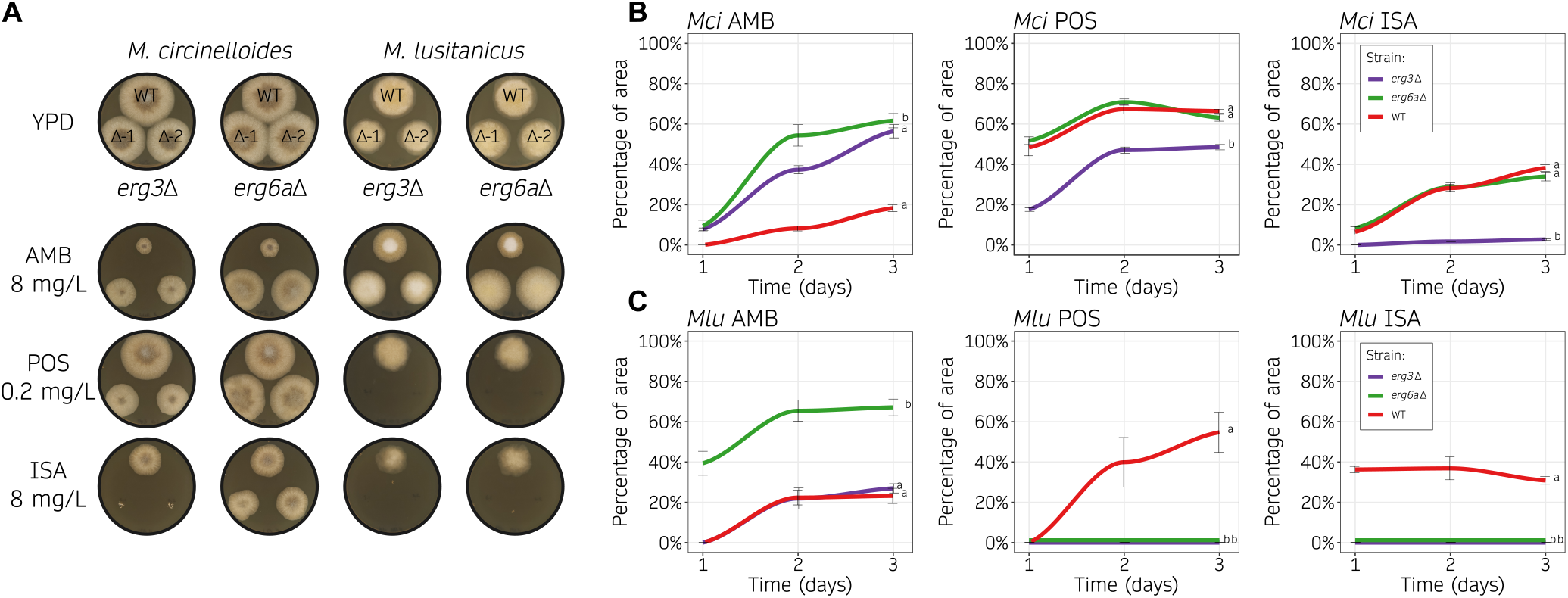
Loss of Erg3 or Erg6a function leads to changes in azole and polyene susceptibility. **(A, B, C)** Amphotericin B (AMB, 8 mg/L), posaconazole (POS, 0.2 mg/mL), and isavuconazole (ISA, 8 mg/mL) susceptibility testing of wild type and ergosterol mutants on solid YPD media. Each species was assayed at its optimal temperature, 30 °C and 26 °C for *M. circinelloides* (*Mci*) and *M. lusitanicus* (*Mlu*), respectively. In **(A)**, representative images of growth after 48 hours in all the drug containing media and the control medium without drug (YPD). Two independently generated mutants were assayed for each gene deletion (Δ-1 and Δ-2). In **(B, C)**, *Mci* **(B)** and *Mlu* **(C)** growth rates from each mutant across time (24-hour intervals) were determined as the percentage of growth area from the same strain cultured on YPD medium with and without drug. Individual values were determined either in six biological replicates for the wild type or in three biological replicates from two independently generated mutants (six total values, grouped together for simplicity as no significant differences were detected), and used to plot a smoothed curve and SD values as black lines. Strains were grouped by letters showing significant growth rate differences across the whole time-course (One-way ANOVA and Tukey HSD test, *p-value* ≤ 0.01).

### Ergosterol profile alteration as the underlying mechanism of antifungal drug resistance

The C-5 desaturase Erg3 and C-24 methyltransferase Erg6 are indispensable components of the ergosterol biosynthetic pathway, responsible for converting lanosterol into ergosterol through a series of catalytic reactions performed by these and other Erg enzymes. Loss-of-function of either enzyme has been shown to alter the ergosterol and related sterol composition in other organisms (42, 43), and we hypothesized that deletion of the *erg3* or *erg6a* genes in *Mucor* species might similarly modify their sterol profiles, explaining the observed changes in antifungal drug susceptibility. To test this hypothesis, we analyzed the sterol profiles of *M. circinelloides* wildtype control, *erg3*Δ, and *erg6a*Δ strains by gas chromatography and mass spectrometry (Figure 5A and Data set S1). Sterols were classified into different categories: C-5(6)-desaturated ergosta-type sterols, C-5(6)-saturated ergosta-type sterols, C-14-methylated sterols, and cholesta-type sterols. This classification aims to simplify sterol characterization and emphasize the accumulation of sterols that may result from Erg3 or Erg6 loss-of-function, as well as exposure to polyene or azole treatments. It is important to note that several sterols classified as cholesta-type may also exhibit C-5(6) desaturation or C-14 methylation.

**Figure 5.**
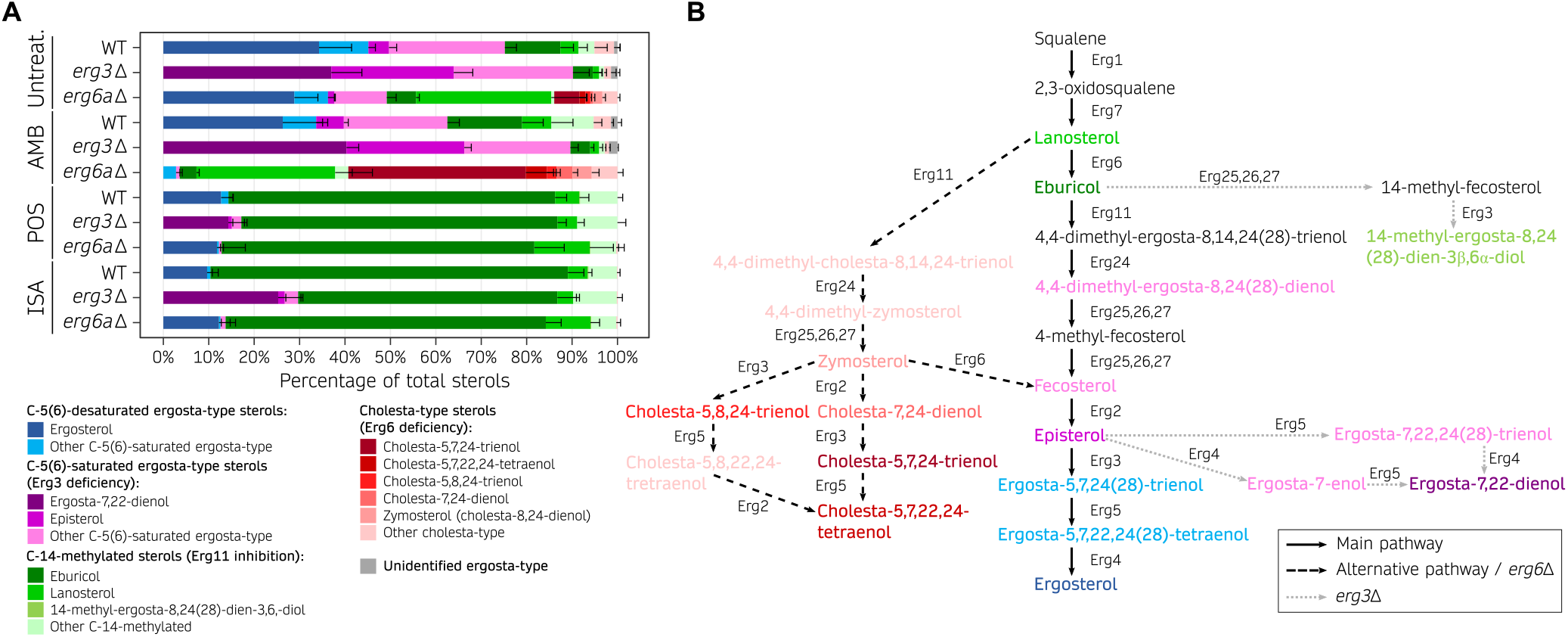
Ergosterol composition upon exposure to first-line mucormycosis treatments is altered by Erg3 and Erg6a deficiencies. **(A)** Percentage of total sterols in *M. circinelloides* wildtype (WT), *erg3*Δ, and *erg6a*Δ strains when exposed to different antifungal treatments (<u>amp</u>hotericin <u>B</u>, <u>pos</u>aconazole, and <u>isa</u>vuconazole) and <u>unt</u>reated cultures, represented in a stacked bar plot. Mean and SD (only upper bound is shown) values were determined from 6 replicates in the WT strain; for each mutation and condition, two independently generated mutants were tested in triplicate (total of 6 replicates). Sterols are classified in four main categories that reflect the expected end products from the main ergosterol pathway [C-5(6)-desaturated sterols], Erg3 deficiency [C-5(6)-saturated sterols], Erg6 deficiency (cholesta-type sterols), and Erg11 inhibition due to azole exposure (14-methylated sterols). Within each category, the most abundant sterols are color-coded to facilitate visualization. **(B)** Model of the ergosterol biosynthetic pathway in Mucorales. The main pathway is depicted by straight, black arrows. Alternative pathways that are relevant in *erg3* (dashed, black arrows) or *erg6a* deletions (dotted, gray arrows) are also shown. For better illustration, sterol compounds are classified and color-coded as in **(A)**.

First, we focused on the sterol profile of *M. circinelloides* cultured in untreated conditions, beginning with the wildtype strain and then the *erg3*Δ and *erg6a*Δ mutants. The relative ergosterol content was lower in *M. circinelloides* wild type (34.3 ± 7.2 %) than in other fungi (19, 21, 44, 45), yet it remains the most abundant sterol (Figure 5A and Data set S1). Intriguingly, we found relatively high levels of C-5(6)-saturated sterols [fecosterol, episterol, ergosta-7,22,24(28)-trienol, and others], as well as C-14-methylated sterols (lanosterol, eburicol, and others). These results indicate that other ergosta-type sterols may act as essential components of *M. circinelloides* membranes in addition to ergosterol. *M. circinelloides erg3* deletion resulted in a *>* 90 % increase in C-5(6)-saturated sterols; notably, these include elevated levels of ergosta-7,22-dienol (37.0 ± 6.7 %), which was not detected in the wildtype strain. More importantly, *erg3*Δ mutation led to complete depletion of C-5(6)-desaturated ergosta-type sterols, including ergosterol, confirming a complete loss of Erg3 function. On the other hand, *erg6a* deletion caused a decrease in ergosterol and an increase in C-24-non-methylated sterols (*>* 40 % lanosterol and cholesta-type sterols), confirming Erg6a activity as a C-24 methyltransferase and suggesting a partial loss of Erg6 function. Surprisingly, *M. circinelloides erg6a*Δ mutants still synthesize C-24-methylated sterols, such as ergosterol, which suggests a compensatory upregulation of additional *erg6* genes. Following this hypothesis, RT-qPCR transcription analysis revealed that upon *erg6a* deletion, *erg6b* is upregulated in regular growth conditions, while *erg6c* is induced upon exposure to amphotericin B (Figure S10B). Overall, these findings are consistent with Erg6a serving as the primary Erg6 enzyme, as Erg6a loss-of-function significantly redirects the ergosterol biosynthetic pathway towards the production of cholesta-type sterols, which is indicative of a severe Erg6 deficiency (Figure 5B, Figure S9, and Figure S10A,C). Erg6 deficiency caused by *erg6a* deletion may be partially compensated by *erg6b* and/or *erg6c* upregulation (Figure S10B), though not completely. The relative abundance of eburicol and absence of zymosterol in the wild-type strain, together with higher relative amounts of lanosterol compared to zymosterol in *erg6a*Δ mutants, evidence that Erg6 precedes Erg11 activity on lanosterol, delimiting the preferred ergosterol biosynthetic pathway in *M. circinelloides* (Figure 5B and Figure S9, center lane).

Next, we analyzed the sterol profile of the wildtype and mutant strains after amphotericin B treatment. Polyene, such as amphotericin B, binding to ergosterol leads to fungal cell death, and is usually correlated with a drastic decrease in ergosterol (10, 11). Indeed, the normalized amount of ergosterol was considerably lower in every strain treated with amphotericin B (Figure S11). The relative sterol contents of amphotericin B-treated *M. circinelloides* wildtype and *erg3*Δ mutant strains were similar to their untreated cultures, except for a marked decrease in ergosterol in the wild type. On the other hand, the relative sterol profile of the *erg6a*Δ mutant was substantially altered by amphotericin B treatment, resulting in a total absence of ergosterol production, abolishment of most C-5(6)-saturated sterols except for fecosterol, which exhibited a drastic decrease, and more importantly, a near 60 % relative abundance of cholesta-5,7,24-trienol and other cholesta-type sterols (Figure 5A and Data set S1). *M. circinelloides erg6a*Δ mutants exhibit an increased resistance to amphotericin B, suggesting that cholesta-type sterol-based membranes are sufficiently stable and that producing these sterols as an alternative to ergosterol may protect the fungus against amphotericin B treatment.

Finally, the sterol content in these strains was also assessed after exposure to two different azoles: posaconazole and isavuconazole. These drugs are the only azoles recommended for salvage therapy against mucormycosis. Azoles target the Erg11 enzyme, inhibiting the crucial C-14 sterol demethylase activity responsible for converting lanosterol or eburicol into the subsequent C-14-demethylated sterol intermediate, ultimately leading to ergosterol (Figure 5B, Figure S9, and Figure S10C). In many fungal species, this blockade in the ergosterol biosynthetic pathways redirects the sterol flow into a different pathway that leads to the accumulation of the toxic sterol 14-methyl-ergosta-8,24(28)-dien-3,6-diol, hereinafter toxic 3,6-diol, which is the cause of the azole antifungal effect. Indeed, the sterol profile revealed that exposure to either azole drug led to the overaccumulation of C-14-methylated sterols, mainly eburicol (*≥* 70 %); and a substantial decrease in ergosterol and other ergosta-type sterols (Figure 5A and Data set S1). But surprisingly, azole treatment did not result in an overaccumulation of the toxic 3,6-diol or its precursor 14-methyl-fecosterol, both frequently found in large proportions in other fungal species (12). Only trace amounts of the toxic 3,6-diol (0.2 ± 0.1 %) were found in the wildtype strain during exposure to both azoles, while *erg3*Δ mutants did not produce the toxic 3,6-diol at all (Figure 5A and Data set S1). These findings reveal that although *M. circinelloides* Erg3 is capable of catalyzing this step, there is a limited utilization of the pathway responsible for converting eburicol into toxic sterols during azole exposure (Figure S10C,D), and are particularly intriguing given the susceptibility of *M. circinelloides* to posaconazole and isavuconazole. Notably, the hypersusceptibility of the *erg3*Δ mutants to these azoles implies that the antifungal effect results from the absence of ergosterol and other stable sterols in the cellular membranes (Figure 5A and Data set S1).

Taken together, our findings define the preferred ergosterol biosynthetic pathway in *M. circinelloides* (Figure 5B and Figure S9, center lane) and reveal that Erg3 and Erg6a play critical roles in antifungal drug susceptibility. The *erg6a* deletion neither impairs growth nor abolishes virulence completely, but results in decreased Erg6 activity and therefore, an abundance of stable cholesta-type sterols over ergosterol during exposure to amphotericin B that accounts for the antifungal resistance observed in *erg6a*Δ mutants. In addition, despite *erg3*Δ mutation leading to increased susceptibility to azoles, we could not detect an abundance of toxic 3,6-diol during azole exposure, suggesting that lacking critical sterols for proper membrane function is driving posaconazole and isavuconazole antifungal effect in Mucorales.

## Discussion

Mucormycoses are fungal infections that present with various clinical manifestations, respond differently to therapy, and result in varying outcomes. These infections are as diverse as the distinct species responsible for causing the disease. Indeed, mucoralean fungi display interspecies differences in antifungal drug susceptibility (2, 46) and interactions with host immunity (47), which can explain disease heterogeneity, and these considerations emphasize the importance of specific pathogenicity models in mucormycosis research. However, these species remain understudied with limited genetic models available to study pathogenesis and therapeutics. Recent taxonomic advances revealed that many *M. circinelloides* actually belong to different phylogenetic species within the *M. circinelloides* species complex (27), with *M. circinelloides* being the most prevalent in mucormycosis patients. However, research on pathogenicity mechanisms have been conducted with *M. lusitanicus*, a sister species previously classified as *M. circinelloides* f. *lusitanicus*, for which no clinical isolates have been found to date (27). This work addresses these limitations by developing a new molecular biology model based on the pathogenic human isolate *M. circinelloides* strain 1006PhL. As a result of this study, this strain that exhibits full virulence in our murine model is now amenable to genetic transformation and mortality rates and fungal burden are consistently high and reproducible, partially due to this pathogenic isolate thermotolerance to the range of mammalian physiological temperatures. This thermotolerance also allows for standardized antifungal drug susceptibility testing at recommended temperatures. Our research has demonstrated that loss-of-function mutations of key target genes, such as *pyrF*, results in a completely avirulent phenotype, which can be reversed by genetic complementation via genomic integration. The necessity of the pyrimidine biosynthetic pathway for virulence is in agreement with research in other fungal model organisms (36, 48–50), demonstrating that this model is suitable for virulence factor characterization and establishing a foundation for novel molecular pathogenesis research to follow.

The only available antifungal agents effective against mucormycosis, namely azoles and liposomal amphotericin B, modify the ergosterol content, emphasizing the crucial role of ergosterol biosynthesis in antifungal drug susceptibility. For instance, gain-of-function mutations affecting Erg11, the target of azole drugs, typically confer azole drug resistance in many fungal pathogens (51). Similarly, Mucorales exhibit intrinsic resistance to voriconazole and fluconazole due to an Erg11 duplication with an amino acid substitution affecting the azole binding site (6). Mucorales can develop antifungal drug resistance not only through classical genetic point mutations (6) and insertional mutations (52), but also via epigenetic modifications that lead to transient loss of drug target function (8, 52, 53). Our findings elucidate the preferred ergosterol biosynthetic pathway in *M. circinelloides*, indicating that C-24 methylation catalyzed by Erg6a occurs prior to C-14 demethylation by Erg11. This result is consistent with our understanding of ergosterol biosynthesis in other filamentous fungi (45), as well as in *Candida* species (21, 42, 54) and *Cryptococcus neoformans* (55), which deviates from the ergosterol pathway proposed in *Saccharomyces cerevisiae* (56). However, the relative content of ergosterol in *M. circinelloides* is reduced compared to these other fungal species, while other sterols such as episterol, lanosterol, and eburicol are more abundant. There is a common misconception that ergosterol is the sole sterol found in all fungi and is essential for fungal membrane stability. While it may be the dominant sterol in many of the Dikarya fungi, there is substantial evidence of alternative sterols serving the same role, especially early-diverging fungi (EDF) (57). Most EDF lack Erg5 and even Erg11, and instead harbor a different set of sterol biosynthetic enzymes, such as 7-dehydrocholesterol reductase (DHCR7), resulting in a wider diversity of sterols and a complete absence of ergosterol (57, 58). While this is not the case in most Mucoromycota species, our research reveals that they possess at least two Erg6 paralogs, and *Mucor* species harbor three: Erg6a, Erg6b, and Erg6c. These additional Erg6 paralogs, which are putative C-24 methyltransferases, may be responsible for *M. circinelloides* unique sterol composition, particularly the abundance of C-24-methylated sterols. As a sister clade to the Dikarya, Mucoromycota represents their closest EDF relatives and therefore, displays a hybrid sterol composition profile between more phylogenetically distant EDF and well-studied Dikarya. Taken together, our findings provide new insights into the evolution of membrane sterol composition in fungi, showing evidence of alternative sterol pathways.

Dissecting the role of the C-5 sterol desaturase Erg3 in the ergosterol pathway of *M. circinelloides* uncovered unexpected peculiarities. Despite a total absence of ergosterol in the *erg3*Δ mutants, our findings indicate that *Mucor* species are able to survive on alternative sterol intermediates, as has been previously reported in *C. glabrata erg11* mutants (59), and similarly to EDF species naturally lacking ergosterol (57, 58). The altered sterol composition may explain the growth delay observed in *erg3*Δ mutants at optimal temperatures, which becomes more pronounced at higher temperatures, as well as the mutants’ complete loss of virulence. Reasonably, the changes in sterol content, possibly accompanied by compensating alterations in other membrane lipids, can cause an unstable equilibrium that enables growth in non-stressful conditions but collapses upon harsh environmental changes, particularly during thermal stress and host interactions. These findings align with the observed *erg3* loss-of-function mutations in *Candida* species, which exhibit increased susceptibility to physiological stresses (60, 61) and reduced hyphal growth under certain conditions (62). Consequently, many of these *erg3* null mutants of *Candida* species have been associated with reduced pathogenicity in animal models, albeit these azole resistant, *erg3* mutant isolates originated from human patients (61–63), and some compensatory mutations can restore their pathogenic potential (64). In addition to these defects, *erg3* loss-of-function mutations confer azole resistance in nearly every *Candida* species (13, 54, 60, 61, 63–67), which might be the reason they are so frequently identified from clinical isolates. Azoles block ergosterol synthesis by inhibiting the C-14 sterol demethylase Erg11, causing an accumulation of Erg11 substrates: lanosterol and eburicol. Typically, these substrates are redirected to a different pathway characterized by the production of 14-methyl-fecosterol (through the subsequent action of Erg25, Erg26, and Erg27), which is then turned into the toxic 3,6-diol through Erg3 C-6 hydroxylase activity. The absence of Erg3 activity prevents the formation of the toxic 3,6-diol and allows accumulation of 14-methyl-fecosterol, accounting for the observed resistance in the *Candida erg3* null mutants (42, 68). Because the loss of Erg3 function prevents ergosterol synthesis in *Candida* species, polyenes are not able to bind the cell membrane nor exert their antifungal effects. As a result, these mutants often exhibit resistance not only to azoles but also to polyenes such as amphotericin B (15, 54, 69, 70). *M. circinelloides erg3*Δ mutants are also more resistant to amphotericin B and, as in *Candida* species, this is likely due to a complete absence of ergosterol that prevents polyene binding. However, these mutants are hypersusceptible to azole drugs instead of more resistant, and neither the wildtype nor the mutant strains produce the toxic 3,6-diol. This finding suggests that Erg3 may exhibit a limited capability for C-6 hydroxylation of C-14-methylated sterols compared to other fungi. Nevertheless, the overaccumulation of eburicol in *M. circinelloides erg3*Δ mutants during azole exposure, together with absence of 14-methyl-fecosterol and 4,14-dimethyl-cholesta-8,24-dienol, imply that the Erg25, Erg26, and Erg27 enzymes may prefer different substrates other than eburicol or are simply unable to initiate C-4 demethylation on C-14-methylated sterols as more likely scenarios. This sterol composition with abundant eburicol and absence of 14-methyl-fecosterol or ergosterol could hinder growth and cell membrane function, explaining the hypersusceptibility to azoles in *M. circinelloides erg3*Δ mutants and the differences compared to *Candida* species.

This study further investigates Erg6, a C-24 methyltrans-ferase that catalyzes an essential step in the biosynthesis of ergosterol. Erg6 function has been studied in several fungal pathogens and revealed as important for growth and development under various conditions, adaptation to cell wall stresses, virulence (17, 19–21, 23), and in some instances, essential for cell viability (45). Despite these defects, the absence of Erg6 activity can provide a selective advantage by conferring amphotericin B drug resistance, as seen in multiple *Candida* species and *Cryptococcus neoformans* (17– 21, 23). Indeed, a recently conducted large-scale study, involving a collection of over four hundred *C. auris* strains evolved from clinical isolates, reported that mutations in *erg6* were the most frequent cause of amphotericin B resistance (71). Exploring the function of Erg6 in *Mucor* species presented additional challenges due to the presence of three different Erg6 paralogs. Our phylogenetic and transcriptomic analyses identified Erg6a as the primary Erg6 enzyme involved in *Mucor* species ergosterol biosynthesis. This hypothesis was confirmed by a substantial reduction in C-24methylated sterols in the *erg6a*Δ mutants, although some residual Erg6 activity persisted. Erg6b and Erg6c are upregulated upon loss of Erg6a function and therefore, might serve as functionally redundant C-24 methyltransferases, accounting for the residual Erg6 activity. As in other fungal species, *Mucor* species *erg6a*Δ mutants exhibit defects in asexual sporulation, a significant delay in virulence, and decreased host dissemination. More importantly, *erg6a*Δ mutants are highly resistant to amphotericin B. The sterol profile of *erg6a*Δ mutants under amphotericin B exposure provided insights into the mechanism of drug resistance. It revealed an abundance of cholesta-type sterols, which bear a closer resemblance to cholesterol found in animal cell membranes. A similar sterol profile was observed in amphotericin B-resistant *C. auris* (21) and *C. glabrata* (17) clinical isolates harboring *erg6* mutations, indicating that cholesta-type sterols are a viable alternative to ergosterol.

Our findings raise questions with respect to a recent study of *M. lusitanicus erg6* mutants suggesting the fungus utilizes cholesta-type sterols to produce desmosterol or cholesterol (72), a process requiring DHCR7 activity. Neither their analysis nor ours identified any C7-saturated sterols and although many EDF harbor DHCR7, mucoralean genomes do not encode this enzyme (58). Notably, the authors observed no changes in susceptibility to amphotericin B in their reported *erg6a* mutants, whereas we identified clear resistance to amphotericin B in multiple independently generated *erg6a*Δ mutants of two different *Mucor* species. We attribute these seemingly conflicting results to potential methodological issues in gene deletion verification, as Bauer et al. presented no DNA-based evidence confirming the complete and homokaryotic deletion of *erg6* gene sequences. Our findings, supported by rigorous genetic methodology, define the most plausible pathway for cholesta-type sterol production and their role in amphotericin B resistance in *Mucor erg6a*Δ mutants, corroborated by research in other fungal pathogens. Given the growing susceptible patient population and dearth of available antifungal drugs, mucormycoses pose substantial challenges to global public health. We propose that mutations and epimutations leading to the loss of Erg6a function may provide a selective advantage to Mucorales by conferring amphotericin B resistance. Although *M. circinelloides erg6a*Δ mutants exhibit minor defects in virulence, they can sustain mammalian physiological temperatures and effectively disseminate the infection. It is likely that the residual Erg6 activity observed in the *erg6a*Δ mutants can rescue the most deleterious consequences of loss of Erg6 function, while still conferring amphotericin B resistance due to cholesta-type sterols production. A comparable scenario was recently reported in *C. auris* evolved isolates, where *erg6* mutations conferring amphotericin B resistance emerged in vivo, occasionally accompanied by other mechanisms that compensate for fitness trade-offs (71). Similarly, the loss of Erg6a function in mucoralean pathogens may arise in vivo and lead to mucormycosis infections that do not respond to first-line antifungal therapies, and may challenge recent and exciting developments in polyene antifungal drug discovery (73). Currently, it is complicated to identify resistant clinical isolates because there are no clinical breakpoints defined for antifungal susceptibility testing in Mucorales (74). Therefore, improving current guidelines to identify clinically amphotericin B-resistant isolates from mucormycosis patients should be a priority. Antifungal susceptibility testing may pose a challenge in identifying epimutant clinical isolates due to the transient and unstable nature of epigenetic changes, which can occur but remain undetected. This adds an additional layer of complexity to the identification of resistant clinical isolates but opens new and exciting avenues for future research.

## Materials and methods

### Fungal strains and culturing conditions

*M. lusitanicus* mutant strains generated in this work derive from the double auxotrophic (Ura-, Leu-) MU402 strain (75), using the selectable marker *pyrG* to complement uracil auxotrophy through genetic transformation. The MU636 strain (30) was used as the wildtype, complemented control (*pyrG*^-^::*pyrG*^+^ *leuA*^-^) for phenotypic screening of *M. lusitanicus* mutants. Similarly, *M. circinelloides* mutant strains derive from *M. circinelloides* 1006PhL (34). A uracil auxotroph amenable to genetic transformation, MIN6 (Ura-) was generated in this work. Uracil auxotrophy was complemented by the selectable marker *pyrF*. For phenotypic analysis, the CPA6 strain (*pyrF*^-^::*pyrF*^+^) was used as a control. All the strains generated in this study are listed in Table S3.

Spores were harvested from YPD solid cultures after 4-5 days and counted to perform experiments. The growth rates were quantified by spot inoculating 500 spores onto rich yeast-peptone-dextrose (YPD) medium and minimal yeast-nitrogen-base (YNB) medium at 26 ^°^C, 30 ^°^C, and 37 ^°^C, and analyzed with Fiji (76).

### Isolation of uracil auxotrophic strain MIN6

10^6^ spores of the wildtype 1006PhL strain were spread-plated on YPD-5-FOA (3 g/L) supplemented with uracil (100 mg/L) and uridine (200 mg/L) and screened for resistant isolates after 3 days. Resistant isolates were rechallenged and those exhibiting stable 5-FOA resistance were inoculated onto YNB media with or without uracil and uridine to determine uracil auxotrophy. Genomic DNA from isolates exhibiting uracil auxotrophy was purified using the MasterPure™ Complete DNA and RNA Purification Kit (Lucigen). The promoter, ORF, and terminator containing sequences from *pyrG* and *pyrF* were PCR-amplified and Sanger-sequenced using primers listed in Table S4.

### Protoplast generation and transformation by electroporation

*M. lusitanicus* transformation was performed following previously established protocols (41). For *M. circinelloides* transformation, MIN6 spores were incubated in yeast-peptone-glucose (YPG) medium at 26 ^°^C until germlings appeared. Cell walls were digested with lysing enzymes from *Trichoderma harzianum* (L-1412, Sigma-Aldrich) and chitosanase (C-0794, Sigma-Aldrich) at 30 ^°^C for 1.5 hours (41). Protoplasts were washed, centrifuged, and resuspended in isotonic buffer. After that, protoplasts were mixed with linear double-stranded DNA (3-6 µg), and subjected to exponential decay waveform electroporation (1,000 V, 25 µF, and 400 Ω). Protoplasts were recovered in YPG at 26 ^°^C for 1 hour, and spread-plated on minimal medium with casamino acids for *Mucor* (MMC) pH = 3.2 to select prototroph transformants after 2-5 days incubation in dark conditions.

For gene deletions, we designed linear DNA constructs containing the selectable marker flanked by 1-kb upstream and downstream regions of the target gene. Either *pyrG* or *pyrF* were used as selectable markers for *M. lusitanicus* and *M. circinelloides*, respectively. In addition, a *M. circinelloides* complemented control strain named CPA6 (*pyrF*-Δ59::*pyrF*^+^) was generated by integrating the wildtype *pyrF* marker into MIN6, replacing the *pyrF* mutated allele (*pyrF*-Δ59). Gene deletions as well as homokaryosis were verified by AFLP; first, after two vegetative passages in selective medium MMC, and subsequently, after every additional passage until homokaryosis was achieved.

### Ortholog search and phylogenetic tree inference

Erg3 and Erg6 protein sequences from *Saccharomyces cerevisiae* were used as queries in a PSI-BLAST v2.12.0 search (77) against a proteomic database of selected species (see Table S5 for a comprehensive list) (32–34, 38–40, 78–90). Every match was subjected to a reciprocal BLASTp search against the *S. cerevisiae* proteome. These putative orthologs (Tables S1 and S2) were aligned using MAFFT v7.475 (91), trimmed with TrimAl v1.4.rev15 (92), and phylogenetic trees with 1,000 ultrafast bootstraps and SH-aLRT replicates were inferred by IQ-TREE v2.2.0.3 (93).

### RNA isolation, sequencing, and data analyses

Duplicated YPD liquid cultures were grown for 16 hours at 26 ^°^C and 250 rpm. Total RNA was purified with a QIAGEN miRNeasy Mini Kit. RNA libraries were prepared using Illumina Stranded Total RNA Prep with Ribo-Zero Gold rRNA Removal Kit, and sequenced to obtain 150-bp paired-end reads. In addition to these, *M. lusitanicus* similar and publicly available data were used for gene expression analyses (52). Quality was assessed by FASTQC v0.11.9 and adapters and low-quality reads removed by TrimGalore! V0.6.7. Processed reads were aligned to either *M. circinelloides* 1006PhL (https://fungidb.org/fungidb/app/record/dataset/DS_8b08c1c31d) or *M. lusitanicus* MU402 genome (https://mycocosm.jgi.doe.gov/Muccir1_3/Muccir1_3.info.html) employing STAR v.2.7.10a (94). Coverage files were generated using bamCompare from Deeptools2 v3.5.1 (95) to merge duplicates into a single bigWig file.

### Synteny analysis

Pairwise synteny among closely related Mucoralean species was assessed using the JCVI toolkit MC-Scan pipeline (96). Genomic plots were generated by Deeptools2 pyGenomeTracks v3.7 (97).

### Virulence and fungal burden assays

4-week old BALB/c mice (Charles River) weighing 20 to 25 g were immunosuppressed with cyclophosphamide via intraperitoneal injection (200 mg/kg of body weight), 2 days prior to infection and every 5 days thereafter. Groups of 10 mice were intravenously challenged via retro-orbital injection with 1 x 10^6^ spores from each of the mutant and control strains.

Fungal burden was quantified in five organs (brain, lung, spleen, kidney, and liver) of three infected mice meeting endpoint criteria per group. Organ homogenates were plated onto YPD supplemented with 1 *µ*g/mL of FK506 to induce yeast growth. Colony forming units (CFU) were quantified after 4 days, and normalized per volume plated and organ weight.

### Antifungal drug susceptibility testing

Antifungal drug susceptibility were determined by broth microdilution using the CLSI and EUCAST standard methodology for molds. Minimal inhibitory concentrations (MIC) were evaluated for amphotericin B, posaconazole, and isavuconazole. 10^5^ spores/mL were incubated in Roswell Park Memorial Institute (RPMI) 1640 medium at different drug concentrations and 35 ^°^C for 24 and 48 hours. Wells were evaluated for visible growth or lack therein. Additionally, 500 spores were spot-inoculated onto solid YPD at specific drug concentrations: 8 mg/L of liposomal amphotericin B (Ambisome, Gilead Sciences), 0.2 mg/L of posaconazole (Noxafil, Merck), and 8 mg/L of isavuconazole (Cresemba, Astellas Pharma). The area of inhibited growth was determined as the ratio of growth area in treated compared to untreated plates using Fiji.

### Ergosterol profile quantification

10^4^ spores were cultured at 35 ^°^C for 48 hours and 60 rpm in RPMI medium (untreated) and RPMI supplemented with half the MIC for each corresponding strain for amphotericin B, posaconazole and isavuconazole drugs. Non-saponifiable lipids were extracted from lyophilized mycelia as previously described (12), with cholesterol added as an internal standard. Sterols were derivatized using 0.1 mL *N*,*O*-Bis(trimethylsilyl)trifluoroacetamide and trimethylsilyl chloride [BSTFA and TMCS, (99:1)] and 0.3 mL anhydrous pyridine and heating at 80 ^°^C for 2 hours (98). TMS-derivatized sterols were analyzed using gas chromatography–mass spectrometry (GS/MS) (Thermo 1300 GC coupled to a Thermo ISQ mass spectrometer, Thermo Scientific) and identified with reference to relative retention times, mass ions and fragmentation spectra. GC/MS data files were analyzed using Xcalibur software (Thermo Scientific). Sterol composition was calculated from peak areas, as a mean of 3 replicates per independently generated mutant (6 replicates per gene deletion) or 6 replicates for the wildtype control strain. The relative quantity of sterols present was determined from the peak areas of the sterol and the internal standard and divided by the dry weight of the sample.

### RT-qPCR analysis

Cultures were obtained using the same conditions as for ergosterol profiling. Total RNA was isolated as previously described, and cDNA synthesized using Maxima™ H Minus cDNA Synthesis Kit (Thermo Scientific). qPCRs were prepared with SYBR green PCR master mix (Applied Biosystems) using primers that specifically amplified *erg6b* and *erg6c* paralogs, and the *vma1* gene served as the endogenous control Table S4), in triplicate, and performed in a QuantStudio™ 3 real-time PCR system.

## Data availability

Raw rRNA-depleted RNA-sequencing datasets obtained from the *M. circinelloides* 1006PhL strain are accessible under PRJNA1046487 NCBI’s Sequence Read Archive (SRA) project accession number. RNA-seq datasets derived from the *M. lusitanicus* MU402 strain were similarly generated for a prior study (PRJNA903107) and are publicly available (52).

## Supporting information

Data Set S1

## ACKNOWLEDGEMENTS

We thank Anna Floyd Averette for her constant technical support. We also extend our gratitude to Marcus Hull for providing technical support during the ergosterol profiling process. We commend Dr. Devjanee Swain Lenz and Duke’s Sequencing and Genomic Technologies Core Facility for their assistance, as well as Thomas Milledge and the Duke Computer Cluster team for their computing resources. This study was supported by Astellas Pharma Investigator Sponsored Research ISR005855/CRES-21B01, and NIH/National Institute of Allergy and Infectious Diseases R01 Grant R01 AI170543 both awarded to J.H. The funders had no role in study design, data collection and interpretation, or the decision to submit the work for publication. J.H. is Codirector and Fellow of the CIFAR program Fungal Kingdom: Threats & Opportunities.

