## Supplemental material for "Alternative ergosterol biosynthetic pathways confer antifungal drug resistance in the human pathogens within the *Mucor* species complex"

#### Supplementary information

#### Extended materials and methods

**Fungal strains and culturing conditions.** *M. lusitanicus* mutant strains generated in this work originate from the *M. lusitanicus* CBS277.49 strain. Its derivative, double auxotrophic (Ura-, Leu-) MU402 strain (75) was used as recipient for genetic transformation procedures, using the selectable marker *pyrG* to complement uracil auxotrophy. The MU636 strain (35) was used as the wildtype, complemented control (*pyrG*$^-$::*pyrG*$^+$ *leuA*$^-$) for phenotypic screening of *M. lusitanicus* mutants. Similarly, *M. circinelloides* mutant strains derive from *M. circinelloides* 1006PhL (39). A uracil auxotroph amenable to genetic transformation, MIN6 (Ura-) was generated in this work. Uridine auxotrophy was complemented by the selectable marker *pyrF*. For phenotypic analysis, the CPA6 strain (*pyrF*$^-$::*pyrG*$^+$) was used as a control. All the strains generated in this study are listed in Table S3.

The growth rate of *erg3* and *erg6a* mutants was quantified by spot inoculating 500 spores onto rich yeast-peptone-dextrose (YPD) medium and minimal yeast-nitrogen-base (YNB) medium at 26 °C, 30 °C, and 37 °C. Images were taken every 24 hours for 3 days, and analyzed with Fiji (76) to determine the area of growth.

**Isolation of uracil auxotrophic strain MIN6.** $10^6$ spores of the wildtype 1006PhL strain were spread-plated on YPD medium containing 5-FOA (3 g/L) supplemented with uracil (100 mg/L) and uridine (200 mg/L) and screened for resistant isolates after 3 days. Spores from resistant colonies were inoculated onto the same medium for a complete vegetative cycle, i.e. germination, colony formation, and asexual sporangiophore development (4-6 days). Colonies exhibiting stable 5-FOA resistance were inoculated onto YNB media with or without uracil and uridine as aforementioned to determine uracil auxotrophy. Genomic DNA from isolates that exhibited uracil auxotrophy was purified using the MasterPure™ Complete DNA and RNA Purification Kit (Lucigen). The promoter, ORF, and terminator containing sequences from *pyrG* and *pyrF* were PCR-amplified and Sanger-sequenced using primers listed in Table S4.

**Protoplast generation and genetic transformation by electroporation.** Protoplast generation and transformation by electroporation of *M. lusitanicus* was performed following previously established protocols (48). For *M. circinelloides* transformation, batches of 1.25 x $10^8$ MIN6 spores were incubated in 25 mL of yeast-peptone-glucose (YPG) medium at 26 °C for 4 to 4.5 hours, specifically until germlings appeared. Cell walls were digested with a 0.01 M sodium phosphate buffer and 0.5 M sorbitol solution (48) containing 5 mg of lysing enzymes from *Trichoderma harzianum* (L-1412, Sigma-Aldrich) and 0.3 µL (15 U/mg) of chitosanase (C-0794, Sigma-Aldrich) at 30 °C for 1.5 hours, ensuring gentle shaking to avoid protoplast lethality. Protoplasts were washed with cold 0.5 M sorbitol solution, centrifuged at low speed ($\leq$ 1,000 rpm), and resuspended in 800 µL of 0.5 M sorbitol. After that, 200 µl of protoplasts were distributed into electroporation cuvettes, mixed with linear DNA (3-6 µg) and subjected to exponential decay waveform electroporation. Electroporation parameters were set to a field strength of 1,000 V ($E = 5kv \cdot cm^{-1}$ ), capacitance of 25 µF, and constant resistance of 400 Ω. Following electroporation, protoplasts were recovered in 1 mL of YPG medium, incubated at 26 °C for 1 hour, and spread-plated on minimal medium with casamino acids for *Mucor* (MMC) pH = 3.2 to select prototroph transformants after 2-5 days incubation in dark conditions.

For the deletion of *erg* genes in *M. lusitanicus* we designed a linear DNA construct containing the selectable marker *pyrG* flanked by 1-kb upstream and downstream regions of either the *erg3* or *erg6a* gene. For *M. circinelloides*, the linear DNA construct was similar but contained the selectable marker *pyrF*. In addition, a linear DNA fragment containing the wildtype *pyrF* marker sequence was used to transform MIN6 (*pyrF*-Δ59) and replace the *pyrF* mutated allele. This allowed us to 1) determine transformation efficiencies at a range of voltages and select the optimum to conduct subsequent electroporation experiments, and 2) generate a complemented control named CPA6 (*pyrF*-Δ59::*pyrF*$^+$). Gene replacements or deletions through integration by homologous recombination of the DNA cassette as well as homokaryosis were verified by PCR-amplification using primers that generate discriminatory amplicons (AFLP); first, transformants were analyzed after two vegetative passages in selective medium MMC, and subsequently, after every additional passage until homokaryosis was achieved.

**Ortholog search and phylogenetic tree inference.** Erg3 and Erg6 protein sequences from *Saccharomyces cerevisiae* were used as queries in a PSI-BLAST v2.12.0 search (77) against a proteomic database encompassing 13 mucoralean species known to cause animal or plant disease, the chytrid *Spizellomyces punctatus* as an outgroup, as well as four well-known fungal pathogens that harbored *erg3* and *erg6* single copy genes, *Schizosaccharomyces pombe* and *S. cerevisiae* (see Table S5 for a comprehensive list of species and proteomes) (37–39, 45–47, 78–90). The glomeromycete *Rhizophagus irregularis* Erg6 homologs were also identified to clarify the phylogenetic inference. After that, every match was subjected to a reciprocal BLASTp search against the *S. cerevisiae* proteome, and positive reciprocal hits were retrieved. These putative orthologs (Tables S1 and S2) were aligned using MAFFT v7.475 (91), the resulting alignments were trimmed with TrimAl v1.4.rev15 gappyout method (92), and phylogenetic trees with 1,000 ultrafast bootstraps and SH-aLRT replicates were inferred from these alignment blocks by IQ-TREE v2.2.0.3 (93), detecting the best substitution models automatically.

**RNA isolation, sequencing, and data analysis.** 2.5 x $10^5$ spore/mL YPD cultures were grown for 16 hours at 26 °C and 250 rpm in duplicates. Total RNA was purified with a QIAGEN miRNeasy Mini Kit. rRNA-depleted RNA libraries were prepared using Illumina Stranded Total RNA Prep with Ribo-Zero Gold rRNA Removal Kit and *M. circinelloides* rDNA specific probes, and cDNA sequenced in a NovaSeq 6000 sequencing system to obtain 150-bp paired-end reads. In addition to these, *M. lusitanicus* similar and publicly available rRNA-depleted RNA sequencing reads were used for gene expression analyses (57). FASTQ dataset quality was assessed by FASTQC v0.11.9 and reads processed by TrimGalore! V0.6.7 to remove adapters and low-quality reads. Processed reads were aligned to either *M. circinelloides* 1006PhL (https://fungidb.org/fungidb/app/record/dataset/DS_8b08c1c31d) or *M. lusitanicus* MU402 genome (https://mycocosm.jgi.doe.gov/Muccir1_3/Muccir1_3.info.html) employing STAR v.2.7.10a (94). Alignments were sorted and classified into forward and reverse stranded by Samtools sort, view, and merge according to their flag values as follows: forward alignments included first-pair alignments from the reverse strand (-f 64 and 16) and second-pair alignments excluding those from the reverse strand (-f 128 and -F 16), and reverse alignments included second-pair alignments from the reverse strand (-f 128 and 16) and first-pair alignments excluding those from the reverse strand (-f 64 and -F 16). Then, coverage files were generated using bamCompare from Deeptools2 v3.5.1 (95) to merge duplicates into a single bigWig file.

**Synteny analysis.** Pairwise synteny among closely related mucoralean species was assessed at the amino acid level using the JCVI toolkit MCScan pipeline (96), defining synteny range to a maximum of 30 kilobases and a minimum of 2 anchors per cluster. Genomic plots were generated by Deeptools2 pyGenomeTracks v3.7 (97) and protein synteny was manually annotated as pink Bezier curves between species. Genes were plotted as single blocks, i.e. ignoring intron-exon information to facilitate rendering and visualization.

**Virulence and fungal burden assays.** 4-week old BALB/c mice (Charles River) weighing approximately 20 to 25 g were used as a host model for virulence assays. Mice were immunosuppressed with cyclophosphamide via intraperitoneal injection (200 mg/kg of body weight), 2 days prior to infection and every 5 days thereafter. Groups of 10 mice were intravenously challenged via retro-orbital injection with 1 x $10^6$ spores from each of the mutant and control strains. Survival and disease symptomatology of each group of mice were monitored every 12 hours, and animals meeting end-point criteria were euthanized by $CO_2$ inhalation and a secondary method.

Fungal burden was quantified in three infected mice meeting end-point criteria per group. Animals were dissected to extract five organs (brain, lung, spleen, kidney, and liver), which were homogenized. Organ homogenates were plated on YPD media supplemented with 1 $\mu$g/mL of FK506 to induce yeast growth and were incubated for 4 days. Colony forming units (CFU) were quantified and normalized per volume plated and organ weight.

**Antifungal drug susceptibility testing.** Antifungal drug susceptibility profiles were determined by broth microdilution using the CLSI and EUCAST standard methodology for molds. Briefly, minimal inhibitory concentrations required to inhibit visible growth by eye (MIC) were evaluated for amphotericin B, posaconazole, and isavuconazole. $10^5$ spores/mL were incubated in Roswell Park Memorial Institute (RPMI) 1640 medium with each drug at 35 °C for 24 and 48 hours. Results were read visually and wells evaluated for visible growth or lack therein. Additionally, 500 spores were spot-inoculated onto YPD solid medium containing drugs at defined concentrations: 8 mg/L of liposomal amphotericin B (Ambisome, Gilead Sciences), 0.2 mg/L of posaconazole (Noxafil, Merck), and 8 mg/L of isavuconazole (Cresemba, Astellas Pharma). Images of growth were taken at 24-hour intervals, and analyzed with Fiji to measure the area of growth. The area of inhibited growth was determined as the ratio of growth area in treated compared to untreated plates.

**Ergosterol profile quantification.** Samples were obtained by growing $10^4$ spores of the mutant and wildtype strain in RPMI media (untreated) and RPMI supplemented with half the MIC for each corresponding strain for amphotericin B, posaconazole and isavuconazole drugs. Cultures were grown at 35 °C for 48 hours and 60 rpm. Non-saponifiable lipids were extracted from lyophilized mycelia as previously described (12), with cholesterol added as an internal standard. Sterols were derivatized using 0.1 mL *N,O*-Bis(trimethylsilyl)trifluoroacetamide and trimethylsilyl chloride [BSTFA and TMCS, (99:1)] and 0.3 mL anhydrous pyridine and heating at 80 °C for 2 hours (98). TMS-derivatized sterols were analyzed using gas chromatography–mass spectrometry (GS/MS) (Thermo 1300 GC coupled to a Thermo ISQ mass spectrometer, Thermo Scientific) and identified with reference to relative retention times, mass ions and fragmentation spectra. GC/MS data files were analyzed using Xcalibur software (Thermo Scientific). Sterol composition was calculated from peak areas, as a mean of 3 replicates per independently generated mutant (6 replicates per gene deletion) or 6 replicates for the wildtype strain 1006PhL. The relative quantity of sterols present was determined from the peak areas of the sterol and the internal standard and divided by the dry weight of the sample.

**RT-qPCR analysis.** Cultures were obtained using the same conditions as for ergosterol profiling. Total RNA was isolated as previously described, and cDNA synthesized using Maxima™ H Minus cDNA Synthesis Kit (Thermo Scientific). qPCRs were prepared with SYBR green PCR master mix (Applied Biosystems) using primers that specifically amplified *erg6b* and *erg6c* paralogs, and the vma1 gene served as the endogenous control (Table S4), in triplicate, and performed in a QuantStudio™ 3 real-time PCR system.

**Data availability.** Raw rRNA-depleted RNA-sequencing datasets obtained from the *M. circinelloides* 1006PhL strain will be accessible upon publication under the following NCBI's Sequence Read Archive (SRA) project accession number: PRJNA1046487. RNA-seq datasets derived from the *M. lusitanicus* MU402 strain were similarly generated for a prior study (57) and are publicly available, assigned to the NCBI's SRA project accession number: PRJNA903107.

### Table S1

| Name | Species | JGI Protein ID | FungiDB Gene ID | EnsemblFungi Gene ID |
|---|---|---|---|---|
| Erg3 | *Candida albicans* SC5314 | 59490 | C1_04770C_A | C1_04770C_A |
| Erg3 | *Choanephora cucurbitarum* NRRL2744 | 516289 | | |
| Erg3 | *Cunninghamella echinulata* NRRL1382 | 185651 | | |
| Erg3 | *Cryptococcus neoformans* H99 | 446 | CNAG_00519 | CNAG_00519 |
| Erg3a | *Lichtheimia corymbifera* JMRC:FSU:9682 | 10330 | LCOR_09389.1 | LCOR_09389.1 |
| Erg3b | *Lichtheimia corymbifera* JMRC:FSU:9682 | 4998 | LCOR_04579.1 | LCOR_04579.1 |
| Erg3a | *Lichtheimia ramosa* JMRC:FSU:6197 | 7192 | | LRAMOSA02887 |
| Erg3b | *Lichtheimia ramosa* JMRC:FSU:6197 | 11099 | | LRAMOSA06288 |
| Erg3 | *Malassezia pachydermatis* CBS1879 | 2056 | Malapachy_0912 | Malapachy_0912 |
| Erg3 | *Mucor circinelloides* 1006PhL | 6367 | HMPREF1544_06438 | HMPREF1544_06438 |
| Erg3 | *Mucor lusitanicus* MU402 | 1381623 | | |
| Erg3 | *Mucor lusitanicus* CBS277.49 v2.0 | 156917 | QYA_156917 | MUCCIDRAFT_156917 |
| Erg3 | *Mucor racemosus* UBOCC-A-109155 | 3973 | | |
| Erg3 | *Phycomyces blakesleeanus* NRRL1555 | 178901 | PHYBL_178901 | |
| Erg3a | *Rhizopus azygosporus* CBS357.93 | | | CU097_003763 |
| Erg3b | *Rhizopus azygosporus* CBS357.93 | | | CU097_010180 |
| Erg3a | *Rhizopus delemar* 99-880 | 4689 | RO3G_07367 | RO3G_07367 |
| Erg3b | *Rhizopus delemar* 99-880 | 7668 | RO3G_13407 | RO3G_13407 |
| Erg3 | *Rhizopus microsporus* var. *microsporus* ATCC52813 | 213209 | | RHIMIDRAFT_213209 |
| Erg3 | *Saccharomyces cerevisiae* S288C | 4075 | YLR056W | YLR056W |
| Erg31 | *Schizosaccharomyces pombe* 972h- | 372 | SPAC1687.16c | SPAC1687.16c |
| Erg32 | *Schizosaccharomyces pombe* 972h- | 2860 | SPBC27B12.03c | SPBC27B12.03c |
| Erg3 | *Spizellomyces punctatus* DAOM BR117 | 5830 | SPPG_02338 | SPPG_02338 |
| Erg3 | *Syncephalastrum racemosum* NRRL2496 | 499858 | | |
| Erg3a | *Sporodiniella umbellata* MES 1446 | 444990 | | |
| Erg3b | *Sporodiniella umbellata* MES 1446 | 496186 | | |
| Erg3 | *Ustilago maydis* 521 | 10884 | UMAG_03593 | UMAG_03593 |

**Table S1.** Erg3 amino acid sequences obtained from the genomes of listed fungal species. Identification numbers from different databases [Joint Genome Institute MycoCosm (JGI, https://mycocosm.jgi.doe.gov), Fungi Database (FungiDB, https://fungidb.org) and Ensembl Fungi (https://fungi.ensembl.org] are indicated to facilitate accessibility and reproducibility.

### Table S2

| Name | Species | JGI Protein ID | FungiDB Gene ID | EnsemblFungi Gene ID |
|---|---|---|---|---|
| Erg6 | *Candida albicans* SC5314 | 57503 | C3_02150C_A | C3_02150C_A |
| Erg6a | *Choanephora cucurbitarum* NRRL2744 | 208609 | | |
| Erg6b | *Choanephora cucurbitarum* NRRL2744 | 449520 | | |
| Erg6a | *Cunninghamella echinulata* NRRL1382 | 280554 | | |
| Erg6 | *Cryptococcus neoformans* H99 | 3420 | CNAG_03819 | CNAG_03819 |
| Erg6a | *Lichtheimia corymbifera* JMRC:FSU:9682 | 1250 | LCOR_01192.1 | LCOR_01192.1 |
| Erg6b | *Lichtheimia corymbifera* JMRC:FSU:9682 | 3633 | LCOR_03386.1 | LCOR_03386.1 |
| Erg6d | *Lichtheimia corymbifera* JMRC:FSU:9682 | 3987 | LCOR_03695.1 | LCOR_03695.1 |
| Erg6a | *Lichtheimia ramosa* JMRC:FSU:6197 | 1343 | | LRAMOSA06893 |
| Erg6b | *Lichtheimia ramosa* JMRC:FSU:6197 | 536 | | LRAMOSA00533 |
| Erg6d | *Lichtheimia ramosa* JMRC:FSU:6197 | 2430 | | LRAMOSA07977 |
| Erg6 | *Malassezia pachydermatis* CBS1879 | 2313 | Malapachy_1146 | Malapachy_1146 |
| Erg6a | *Mucor circinelloides* 1006PhL | 10718 | HMPREF1544_10842 | HMPREF1544_10842 |
| Erg6b | *Mucor circinelloides* 1006PhL | 8587 | HMPREF1544_08689 | HMPREF1544_08689 |
| Erg6c | *Mucor circinelloides* 1006PhL | 9624 | HMPREF1544_09736 | HMPREF1544_09736 |
| Erg6a | *Mucor lusitanicus* MU402 | 1319648 | | |
| Erg6b | *Mucor lusitanicus* MU402 | 1325190 | | |
| Erg6c | *Mucor lusitanicus* MU402 | 1448211 | | |
| Erg6a | *Mucor lusitanicus* CBS277.49 v2.0 | 155859 | QYA_155859 | MUCCIDRAFT_155859 |
| Erg6b | *Mucor lusitanicus* CBS277.49 v2.0 | 151310 | QYA_151310 | MUCCIDRAFT_151310 |
| Erg6c | *Mucor lusitanicus* CBS277.49 v2.0 | 74496 | QYA_74496 | MUCCIDRAFT_74496 |
| Erg6a | *Mucor racemosus* UBOCC-A-109155 | 9296 | | |
| Erg6b | *Mucor racemosus* UBOCC-A-109155 | 5625 | | |
| Erg6c | *Mucor racemosus* UBOCC-A-109155 | 11127 | | |
| Erg6a | *Phycomyces blakesleeanus* NRRL1555 | 128748 | PHYBL_128748 | |
| Erg6b | *Phycomyces blakesleeanus* NRRL1555 | 114967 | PHYBL_114967 | |
| Erg6a | *Rhizopus azygosporus* CBS357.93 | | | CU097_001869 |
| Erg6a | *Rhizopus delemar* 99-880 | 9579 | RO3G_15767 | RO3G_15767 |
| Erg6e | *Rhizopus delemar* 99-880 | 1008 | RO3G_16049 | RO3G_16049 |
| Erg6a | *Rhizophagus irregularis* DAOM 197198 | 1517096 | | |
| Erg6b | *Rhizophagus irregularis* DAOM 197198 | 1668347 | | |
| Erg6a | *Rhizopus microsporus* var. *microsporus* ATCC52813 | 203690 | | RHIMIDRAFT_203690 |
| Erg6 | *Saccharomyces cerevisiae* S288C | 4664 | YML008C | YML008C |
| Erg6 | *Schizosaccharomyces pombe* 972h- | 3100 | SPBC16E9.05 | SPBC16E9.05 |
| Erg6 | *Spizellomyces punctatus* DAOM BR117 | 8734 | SPPG_05059 | SPPG_05059 |
| Erg6a | *Syncephalastrum racemosum* NRRL2496 | 499603 | | |
| Erg6b | *Syncephalastrum racemosum* NRRL2496 | 466111 | | |
| Erg6a | *Sporodiniella umbellata* MES 1446 | 459696 | | |
| Erg6e | *Sporodiniella umbellata* MES 1446 | 469068 | | |
| Erg6 | *Ustilago maydis* 521 | 10455 | UMAG_03182 | UMAG_03182 |

**Table S2.** Erg6 amino acid sequences obtained from the genomes of listed fungal species. Identification numbers from different databases [Joint Genome Institute MycoCosm (JGI, https://mycocosm.jgi.doe.gov), Fungi Database (FungiDB, https://fungidb.org) and Ensembl Fungi (https://fungi.ensembl.org] are indicated to facilitate accessibility and reproducibility.

### Table S3

| Strain | Species | Genotype | Reference |
|---|---|---|---|
| MU402 | *M. lusitanicus* | *pyrG*$^-$ *leuA*$^-$ | (75) |
| MU636 | *M. lusitanicus* | *pyrG*$^-$::*pyrG*$^+$ *leuA*$^-$ | (35) |
| MIN11 | *M. lusitanicus* | *erg3*::*pyrG leuA-* | This study |
| MIN12 | *M. lusitanicus* | *erg3*::*pyrG leuA-* | This study |
| MIN13 | *M. lusitanicus* | *erg6a*::*pyrG leuA-* | This study |
| MIN14 | *M. lusitanicus* | *erg6a*::*pyrG leuA-* | This study |
| 1006PhL | *M. circinelloides* | wildtype | (39) |
| MIN6 | *M. circinelloides* | *pyrF*-59Δ | This study |
| CPA6 | *M. circinelloides* | *pyrF*-59Δ::*pyrF*$^+$ | This study |
| MIN15 | *M. circinelloides* | *erg3*::*pyrF* | This study |
| MIN16 | *M. circinelloides* | *erg3*::*pyrF* | This study |
| MIN17 | *M. circinelloides* | *erg6a*::*pyrF* | This study |
| MIN18 | *M. circinelloides* | *erg6a*::*pyrF* | This study |

**Table S3.** Strains used in this study, showing from which species they derive, genotype, and references.

### Table S4

| Name | Species | Sequence (5'⟶3') | Description |
|---|---|---|---|
| JOHE51488 | *Mlu* | TGCCTCAGCATTGGTACTTG | *pyrG* PCR |
| JOHE51489 | *Mlu* | GTACACTGGCCATGCTATCG | *pyrG* PCR |
| JOHE51528 | *Mlu* | CGATAGCATGGCCAGTGTACTAGAGTTTACGATGCAGGCCAGT | *erg3* deletion, overlap with *pyrG* |
| JOHE51539 | *Mlu* | CAAGTACCAATGCTGAGGCAATTTCTGTGTGTCAGTCCGCAC | *erg3* deletion, overlap with *pyrG* |
| JOHE51529 | *Mlu* | ATTAGAACAGAAGGGCAGTCGG | *erg3* deletion |
| JOHE51538 | *Mlu* | CGATCATGCTTACGGTGGTTGAGT | *erg3* deletion |
| JOHE51540 | *Mlu* | CGATAGCATGGCCAGTGTACGTGTGCCAAGTGTAGATGTTGTG | *erg6a* deletion, overlap with *pyrG* |
| JOHE51552 | *Mlu* | CAAGTACCAATGCTGAGGCACATCTGTCTCAATATCCGTCGTC | *erg6a* deletion, overlap with *pyrG* |
| JOHE51541 | *Mlu* | ATTTGCCCGCTGTAGATGATAC | *erg6a* deletion |
| JOHE51551 | *Mlu* | AGTGTAGCAAAAGTTGCCCTTG | *erg6a* deletion |
| JOHE51511 | *Mlu* | ACCTTGAGCACACAAACAAAGG | *pyrG,* 5' junction PCR |
| JOHE51510 | *Mlu* | CCTTTGTTTGTGTGCTCAAGGT | *pyrG,* 3' junction PCR |
| JOHE51533 | *Mlu* | ACAGGAAGTGAGTACAACGGACA | *erg3*, 5' junction PCR |
| JOHE51534 | *Mlu* | CATCGCCTCATACTACTCAAAGC | *erg3*, 3' junction PCR |
| JOHE51531 | *Mlu* | TGTGGTATCTACTGGTTCCATCG | *erg3,* WT allele PCR |
| JOHE51536 | *Mlu* | ATGTTGTGTGTCGAGTAGCACC | *erg3,* WT allele PCR |
| JOHE51545 | *Mlu* | GTCTATGTGGCGCTCAATTCTAC | *erg6a*, 5' junction PCR |
| JOHE51546 | *Mlu* | GCGTCCTCTTCCATTCCTACTAC | *erg6a*, 3' junction PCR |
| JOHE51543 | *Mlu* | GCTGGTGATATTTACGAGTCTGC | *erg6a,* WT allele PCR |
| JOHE51549 | *Mlu* | GGAGAGATTTGATGGTGGTGTT | *erg6a,* WT allele PCR |
| JOHE51498 | *Mci* | AGAATGCCAGACCTGAATTTTTGG | *pyrF* PCR |
| JOHE51499 | *Mci* | AATAGTAATACCCTCTGCCAACGG | *pyrF* PCR |
| JOHE51623 | *Mci* | CCGTTGGCAGAGGGTATTACTATTGATATGGTAGCGAACCCGATG | *erg3* deletion, overlap with *pyrF* |
| JOHE51628 | *Mci* | CCAAAAATTCAGGTCTGGCATTCTAATTTCTGTGTGTCAGTCCGCAC | *erg3* deletion, overlap with *pyrF* |
| JOHE51624 | *Mci* | GGATGAGTCTTCAGGTGCAGTAA | *erg3* deletion |
| JOHE51627 | *Mci* | AATCCGAGAATGATGTGTGAATACC | *erg3* deletion |
| JOHE51630 | *Mci* | CCGTTGGCAGAGGGTATTACTATTCAACATCAAATCTCTCCCGTTC | *erg6a* deletion, overlap with *pyrF* |
| JOHE51634 | *Mci* | CCAAAAATTCAGGTCTGGCATTCTCTCAATATCCGTCTTCCCCTTAC | *erg6a* deletion, overlap with *pyrF* |
| JOHE51629 | *Mci* | AAGATTGGAGCACCTCAAGAGAC | *erg6a* deletion |
| JOHE51633 | *Mci* | AGTCGTGCAGCAGGTAGTTTTAG | *erg6a* deletion |
| JOHE51503 | *Mci* | GAGCAGCAGCATAGAAAGTACCA | *pyrF,* 5' junction PCR |
| JOHE51504 | *Mci* | TATCAAGAAGGACTAGGCTTGCC | *pyrF,* 3' junction PCR |
| JOHE51625 | *Mci* | TGTCACCTACACGAATCTTACCC | *erg3*, 5' junction PCR |
| JOHE51626 | *Mci* | AGAGAATATGGCGCTTCAGAGAC | *erg3*, 3' junction PCR |
| JOHE51625 | *Mci* | TGTCACCTACACGAATCTTACCC | *erg3,* WT allele PCR |
| JOHE51637 | *Mci* | TGTTCATCGGTGGGTTGACG | *erg3,* WT allele PCR |
| JOHE51631 | *Mci* | CTGATTTCACAGTACGCCTGAGT | *erg6a*, 5' junction PCR |
| JOHE51632 | *Mci* | GGTAAAGAATATGGGCACAGGTC | *erg6a*, 3' junction PCR |
| JOHE51631 | *Mci* | CTGATTTCACAGTACGCCTGAGT | *erg6a,* WT allele PCR |
| JOHE51641 | *Mci* | GCATCACCAACATTGGCAAG | *erg6a,* WT allele PCR |
| JOHE51712 | *Mci* | GCAAGGCGATATGGGTGTTG | *erg6b,* qPCR |
| JOHE51713 | *Mci* | GACTGAAGAGCGAAGAAACCC | *erg6b,* qPCR |
| JOHE51714 | *Mci* | ACTCAAGCTGGTCGGTATGC | *erg6c,* qPCR |
| JOHE51715 | *Mci* | ACAACAGAATCGCCTGCAAC | *erg6c,* qPCR |
| JOHE51665 | *Mci* | AGGGTCAATACACTCTTGAAG | *vma1,* qPCR |
| JOHE51667 | *Mci* | GGAGAACGAACAGGCCAGAG | *vma1,* qPCR |

**Table S4.** Species-specific primers [*M. lusitanicus* (*Mlu*) or *M. circinelloides* (*Mci*)] used in this study, showing systematic name, nucleotide sequence, and a brief description explaining its use.

### Table S5

| Species | Database | Reference |
|---|---|---|
| *Candida albicans* SC5314 | JGI | (78) |
| *Choanephora cucurbitarum* NRRL2744 v1.0 | JGI | (79) |
| *Cryptococcus neoformans* var. *grubii* H99 | JGI | (80) |
| *Cunninghamella echinulata* NRRL1382 v1.0 | JGI | (79) |
| *Lichtheimia corymbifera* JMRC:FSU:9682 | JGI | (45) |
| *Lichtheimia ramosa* JMRC:FSU:6197 | JGI | (81) |
| *Malassezia pachydermatis* CBS1879 | JGI | (82) |
| *Mucor circinelloides* 1006PhL | JGI | (39) |
| *Mucor lusitanicus* MU402 v1.0 | JGI | (38) |
| *Mucor racemosus* UBOCC-A-109155 | JGI | (83) |
| *Phycomyces blakesleeanus* NRRL1555 v2.0 | JGI | (37) |
| *Rhizophagus irregularis* DAOM 197198 v2.0 | JGI | (84) |
| *Rhizopus azygosporus* CBS357.93 | EnsemblFungi | (47) |
| *Rhizopus delemar* 99-880 | JGI | (46) |
| *Rhizopus microsporus* var. *microsporus* ATCC52813 v1.0 | JGI | (85) |
| *Saccharomyces cerevisiae* S288C | JGI | (86) |
| *Schizosaccharomyces pombe* 972h- | JGI | (87) |
| *Spizellomyces punctatus* DAOM BR117 | JGI | (88) |
| *Sporodiniella umbellata* MES 1446 v1.0 | JGI | (79) |
| *Syncephalastrum racemosum* NRRL2496 v1.0 | JGI | (89) |
| *Ustilago maydis* 521 | JGI | (90) |

**Table S5.** Proteomes from these species were obtained from publicly available repositories either at the Joint Genome Institute MycoCosm (JGI, https://mycocosm.jgi.doe.gov) or, if unavailable, at the EnsemblFungi (https://fungi.ensembl.org) databases.

### Figure S1

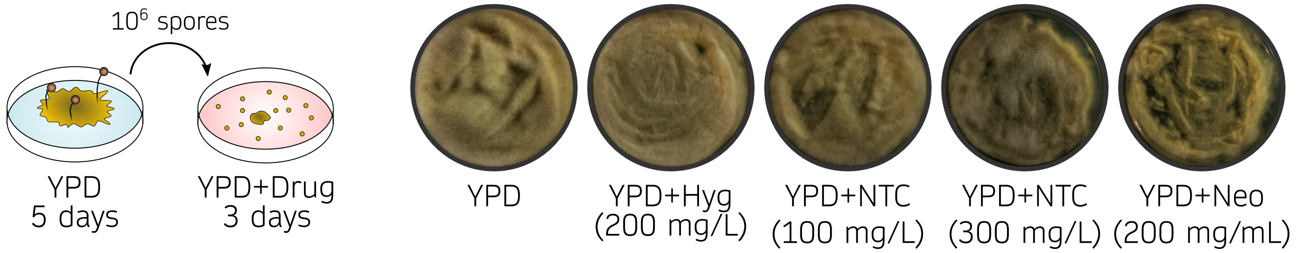


**Figure S1. Evaluation of dominant resistance markers in *M. circinelloides*.** $10^6$ spores from *M. circinelloides* wildtype strain growing for 48 hours on YPD medium supplemented with Hygromycin (Hyg), Nourseothricin (NTC), and Neomycin (Neo) at the indicated concentrations.

### Figure S2

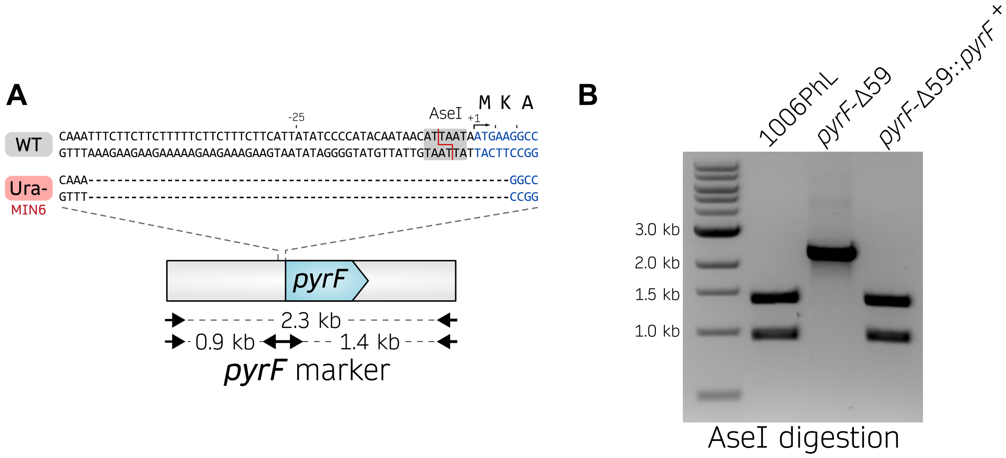


**Figure S2. Complementation of the *pyrF* mutant allele in MIN6. (A)** Wildtype and MIN6 mutant sequences for *pyrF*. The AseI restriction site is indicated in the wildtype 5' region of the *pyrF* gene, while this restriction site is missing in MIN6. **(B)** PCR-amplification followed by AseI digestion confirming *pyrF* complementation.

### Figure S3

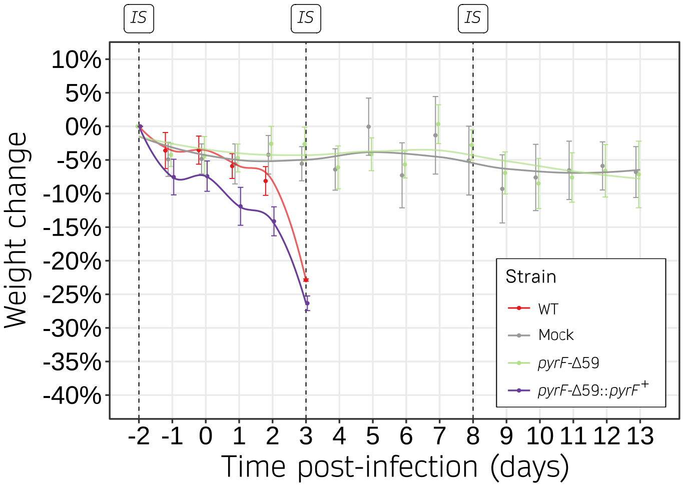


**Figure S3. Weight monitoring of mice infected with uracil auxotrophic and prototrophic strains.** Body weight of animals infected with wildtype 1006PhL, *pyrF*-59Δ auxotrophic strain, and *pyrF*-59Δ::*pyrF*$^{+}$ complemented strain. Animals infected with prototrophic strains (red and purple) show increased weight loss over time compared to animals infected with the auxotrophic strain (green) or PBS (gray). Immunosuppressive (IS) treatments are indicated as dotted lines.

### Figure S4

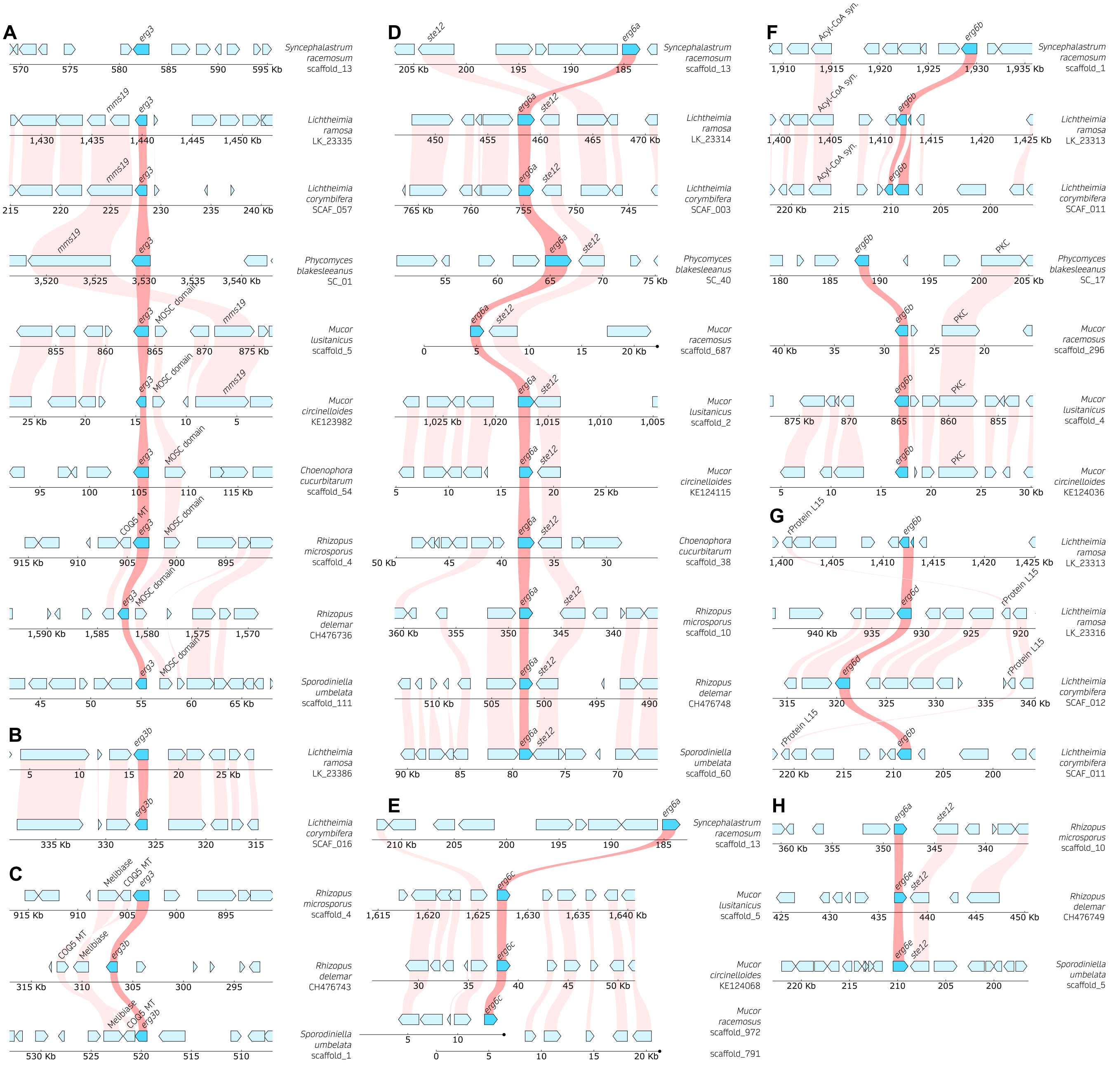


**Figure S4. Synteny of *erg3* and *erg6* orthologs within the Mucoromycota.** Gene synteny of the indicated species and chromosome locations of genes **(A)** *erg3*, **(B)** *erg3b* in *Lichtheimia* spp., **(C)** *erg3b* in *Rhizopus* spp., **(D)** *erg6a*, **(E)** *erg6c*, **(F)** *erg6b*, **(G)** *erg6d*, and **(H)** *erg6e*. Species pairwise comparisons were selected according to closest phylogenetic relationship and when possible, arranged similarly to species in clade *erg6a* from Figure 2B. The genome coordinates are indicated for each genome, and gene annotation is depicted as blue blocks highlighting the *erg* genes in cyan blue. Interspecies synteny among genes is depicted as red shading for *erg* genes and as pink shading for neighboring genes.

### Figure S5

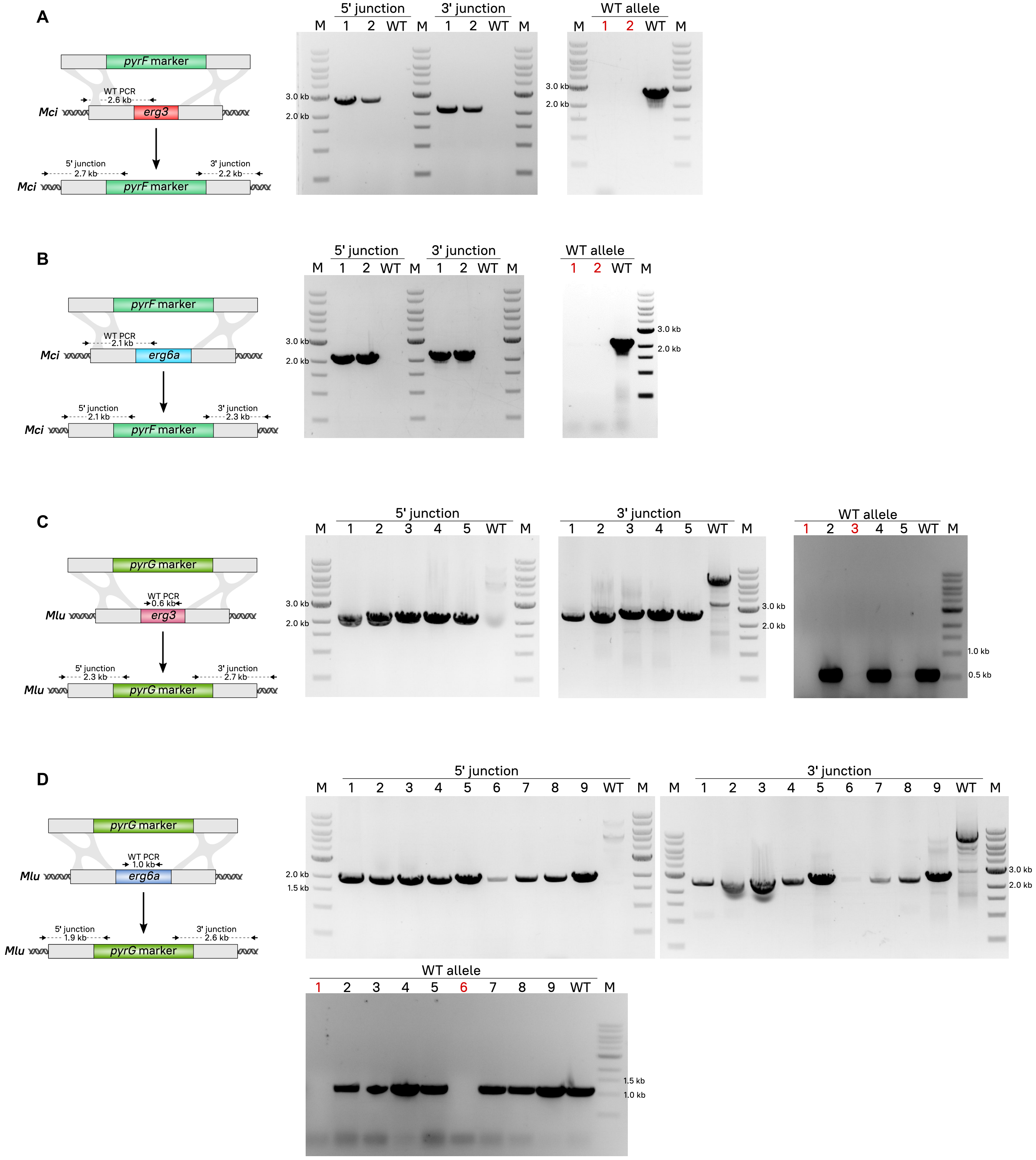


**Figure S5. Generation of mutant strains for *erg3* and *erg6a* genes in *M. circinelloides* and *M. lusitanicus*.** Diagram (left) and AFLP agarose gel images (right) showing the integration of either the *pyrF* or *pyrG* marker replacing the corresponding *erg* gene: deletion of *erg3* **(A)** and *erg6a* **(B)** in *M. circinelloides*; deletion of *erg3* **(C)** and *erg6a* **(D)** in *M. lusitanicus.* Lanes are numbered to indicate transformant isolates, except for M lanes that show a DNA ladder for fragment size comparison. Arrows mark the annealing regions for specific primers used to confirm the integration by two different PCR reactions: 5' junction, and 3' junction. Mutant homokaryosis was confirmed by absence of the wildtype (WT) allele, and homokaryotic mutants generated by independent transformation experiments are highlighted in red.

### Figure S6

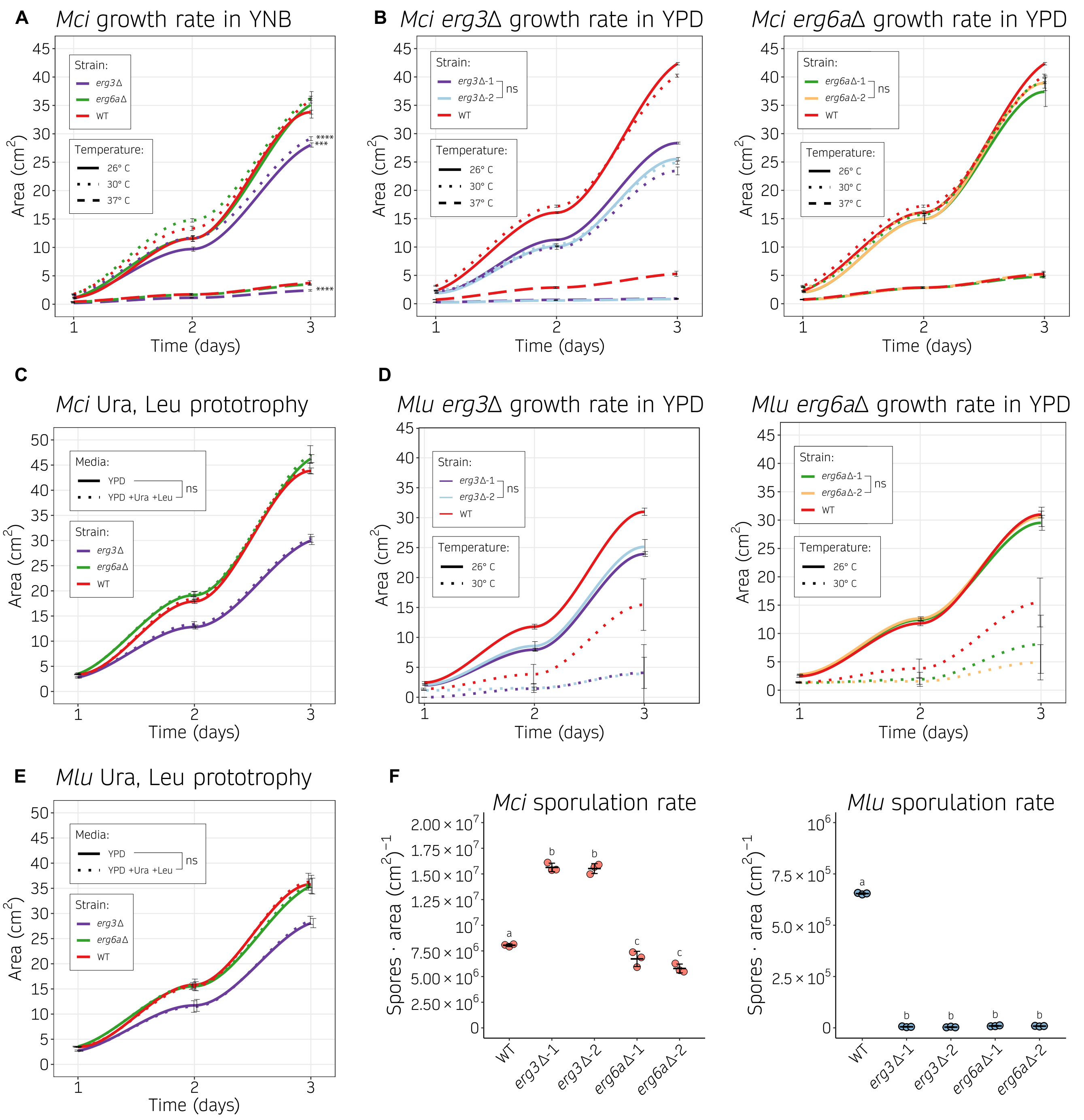


**Figure S6. Growth and sporulation rates of *Mucor* species *erg3*Δ and *erg6a*Δ independently generated mutants. (A-E)** Growth rate of *Mucor* species strains is depicted as color-coded lines, and temperature or media supplementation are indicated as distinct line types. The growth rate was determined as the area of growth after 24-hour periods. Each experiment was done using six replicates for the wildtype strain (WT), or three biological replicates for each pair of independently generated mutants (Δ-1 and Δ-2) accounting for six total replicates for each deletion and condition tested. Values are plotted as a smoothed curve and SD values as black lines. Growth rate was determined in mutant strains from **(A)** *M. circinelloides* growing in minimal medium YNB; **(B)** *M. circinelloides* in rich medium YPD; **(C)** *M. circinelloides* in rich medium YPD with or without uridine and leucine supplementation; **(D)** *M. lusitanicus* in rich medium YPD; and **(E)** *M. lusitanicus* in rich medium YPD with or without uridine and leucine supplementation. One-way ANOVA and Tukey HSD test were conducted for each temperature group and shown as asterisks (** *p-value* $\leq$ 0.01, *** *p-value* $\leq$ 0.001, and **** *p-value* $\leq$ 0.0001) in **(A)**, and to highlight that independently generated mutants harboring the same deletion did not show significant differences (ns) in **(B-D)**, as well as validating the strains prototrophy by showing no significant differences (ns) with uridine and leucine supplementation in **(C-E)**. **(F)** Sporulation rate from *M. circinelloides* or *M. lusitanicus* wildtype strains and two *erg3*Δ or *erg6a*Δ independently generated mutants (Δ-1 and Δ-2) was assessed as the number of spores per $cm^2$ after growing in MMC medium for 48 hours. Strains are grouped by letters, each group showing significant sporulation differences between them (One-way ANOVA and Tukey HSD test).

### Figure S7

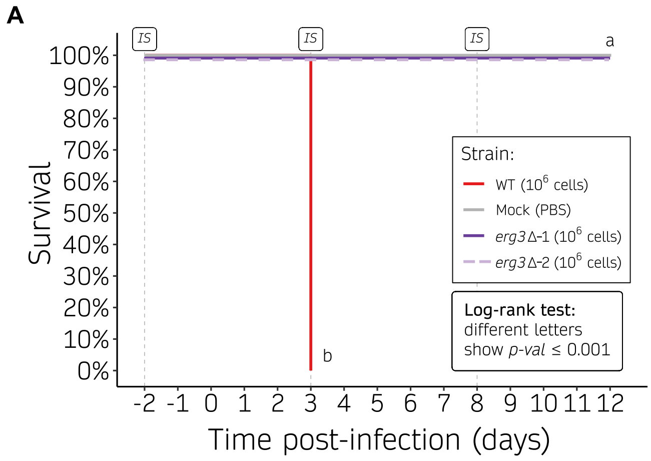


**Figure S7. Survival curves of mice infected with *M. circinelloides erg3*Δ and *erg6a*Δ independently generated mutants.** Kaplan-Meier survival curves of immunosuppressed mice infected with $10^6$ spores from the wildtype strain and either **(A)** *erg3*Δ-1 and *erg3*Δ-2 mutants, or **(B)** *erg6a*Δ-1 and *erg6a*Δ-2. Groups of five mice were inoculated with each independently generated mutant, accounting for ten total mice per deletion or wildtype strain. Strains showing significant differences in virulence assessed by a Log-rank test are indicated as different letters (*p-value* $\leq$ 0.05), confirming that independently generated mutants harboring the same deletion did not show significant differences in virulence. Immunosuppressive treatments are indicated (IS).

### Figure S8

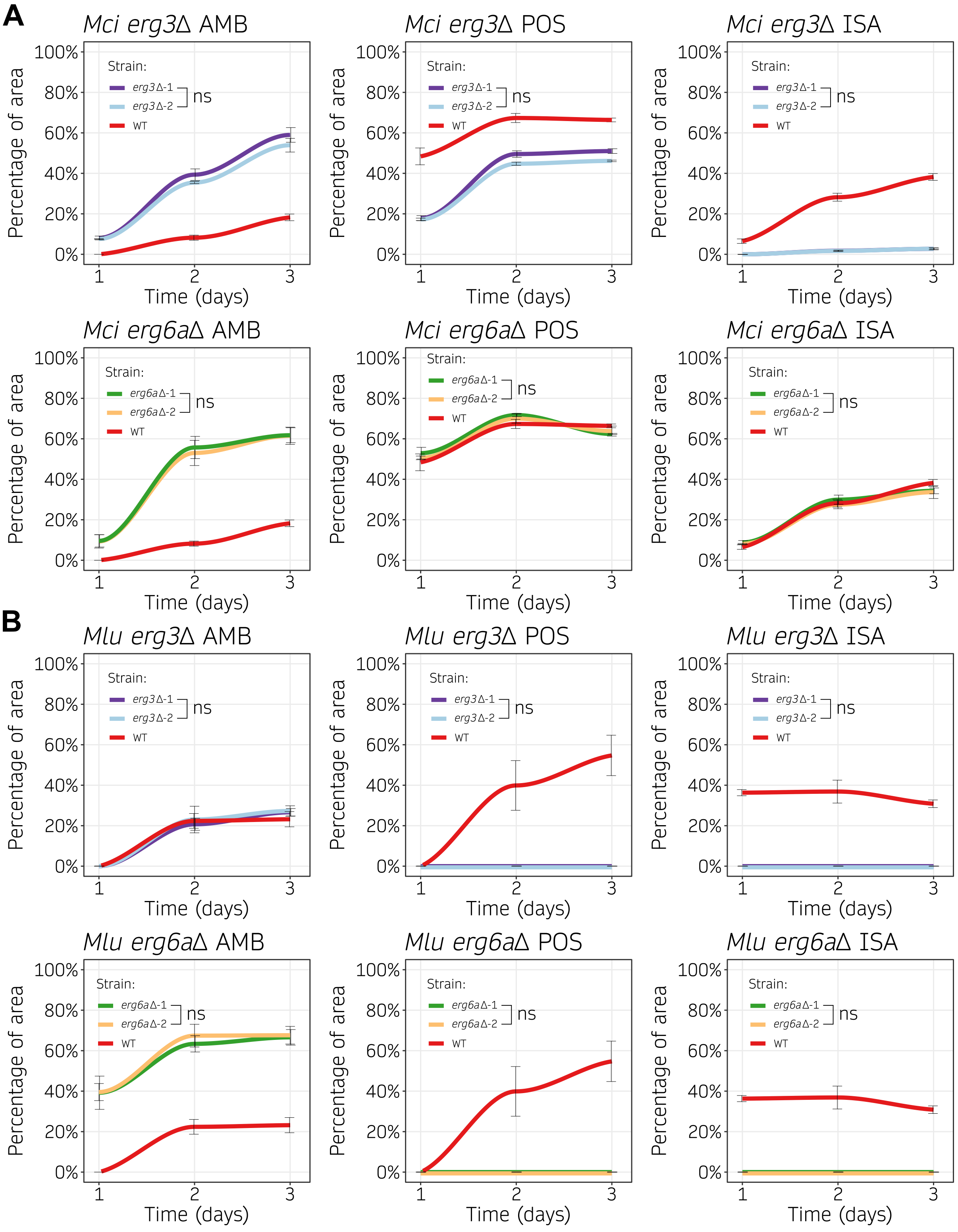


**Figure S8. Growth rate during antifungal exposure in *Mucor* species *erg3*Δ and *erg6a*Δ independently generated mutants.** Growth across time (24-hour intervals) from independently generated *erg3*Δ and *erg6a*Δ mutants (Δ-1 and Δ-2) in medium containing amphotericin B (AMB, 8 mg/L), posaconazole (POS, 0.2 mg/mL), or isavuconazole (ISA, 8 mg/mL) was determined as the percentage of growth area from the same strain cultured in control medium without drug. Each species was assayed at its optimal temperature, 30 °C and 26 °C for *M. circinelloides* (*Mci*) and *M. lusitanicus* (*Mlu*), respectively. Individual values were collected either in six biological replicates for the wild type or in three biological replicates from each of the two independently generated mutants to plot a color-coded smoothed curve and SD values (black lines). Independently generated mutants exhibiting the same deletion did not show significant differences (ns, One-way ANOVA and Tukey HSD test, p-value $\leq$ 0.05).

### Figure S9

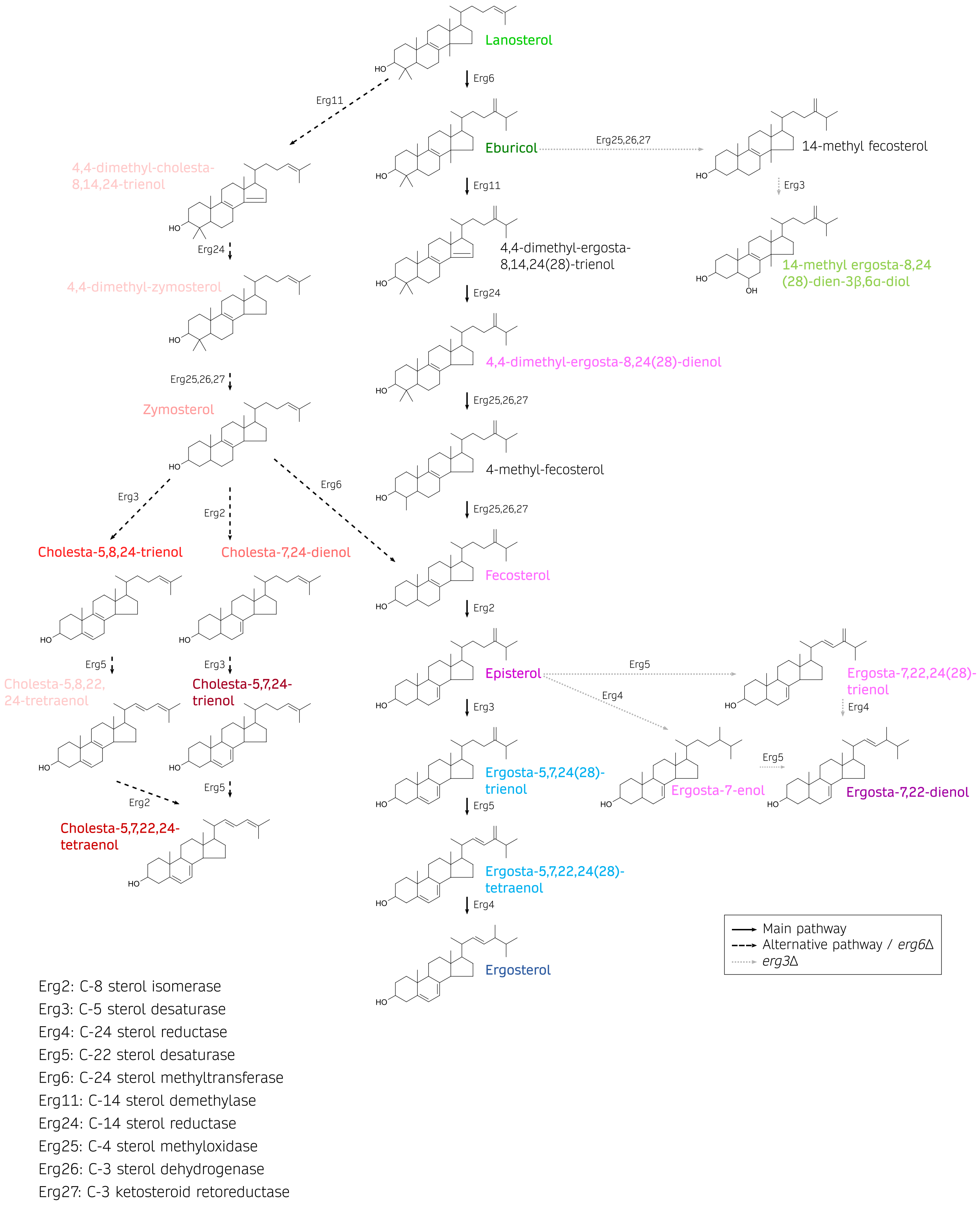


**Figure S9. Model of the ergosterol biosynthetic pathway in Mucorales.** Pathway schematics showing the compounds (line-angle formula and systematic name) and enzymes (enzymatic activity at bottom legend) comprising the ergosterol biosynthetic pathway in Mucorales. The preferred pathway is depicted by straight, black arrows. Alternative pathways that are relevant in *erg3* (dashed, black arrows) or *erg6a* deletions (dotted, gray arrows) are also shown. For better illustration, sterol compounds are classified and color-coded as in Figure 5A.

### Figure S10

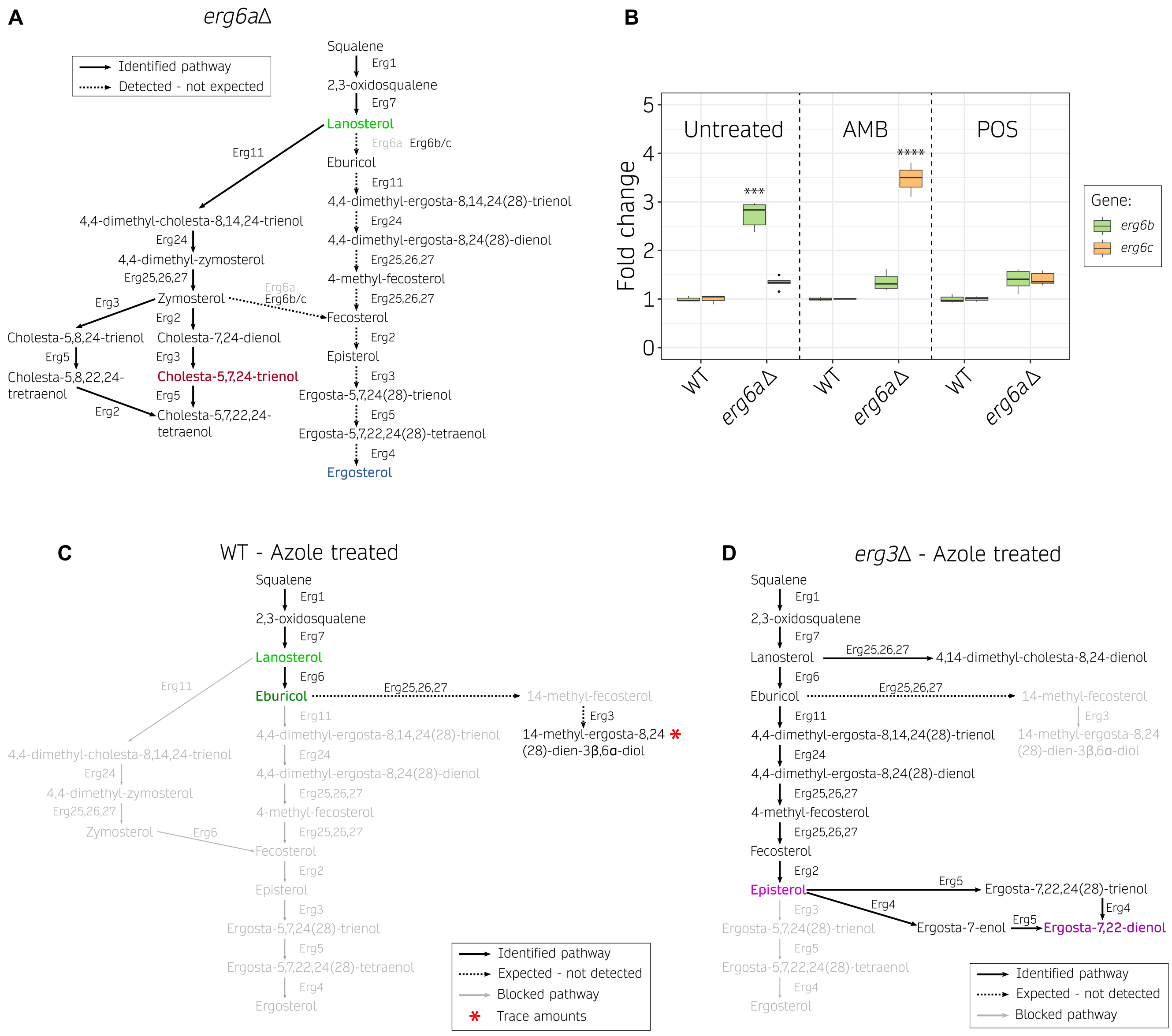


**Figure S10. Alternative sterol biosynthetic pathways during azole and amphotericin B treatment in *M. circinelloides*. (A)** Cholesta-type sterol accumulation in the *erg6a*Δ mutant with amphotericin B treatment. Erg6a loss-of-function is indicated with gray color, and leads to the accumulation of identified cholesta-type sterols defined by straight, black arrows. Unexpected, residual Erg6 activity was detected by the accumulation of C-24 methylated sterols defined by dashed, black arrows. These sterols, including ergosterol, indicate residual Erg6 activity possibly due to additional Erg6 paralogs, Erg6b or Erg6c. **(B)** Box plot of *erg6b* and *erg6c* transcription differences due to *erg6a*Δ mutation compared to the wildtype strain in untreated conditions and upon amphotericin B (AMB) and posaconazole (POS) exposure at each strain half MIC values. Fold-change differential expression levels were quantified by RT-qPCR and normalized to *vma1*, a vacuolar ATPase encoding gene that is constitutively expressed, as an internal control. Minimum, maximum, quartile, and median values correspond to technical triplicates of wildtype, *erg6a*Δ-1, and *erg6a*Δ-2 strains; *erg6a*Δ mutant strain values were pooled together into a total of 6 replicates after no significant differences were observed between them. Significant differences are denoted by asterisks (*** for P $\leq$ 0.005 and **** for P $\leq$ 0.0001 in a One-way ANOVA and Tukey HSD test).**(C, D)** Accumulation of 14-methylated sterols and other ergosta-type sterols during posaconazole and isavuconazole (azoles) exposure. Identified compounds and reactions are indicated with black arrows, while unidentified compounds and reactions are shown as gray arrows (blocked pathway). Blockades due to azole-mediated Erg11 inhibition in the wildtype strain **(B)** and in *erg3*Δ mutants **(C)** are depicted. Dashed, black arrows indicate the expected accumulation of 14-methyl-fecosterol and the toxic 3,6-diol, but only trace amounts of the toxic 3,6-diol were detected in the wild-type strain during azole treatment. The most abundant sterol compounds in each pathway are color-coded as follows: green (14-methylated sterols), pink (C-5(6)-saturated ergosta-type sterols), red (cholesta-type sterols), and blue (C-5(6)-desaturated ergosta-type sterols).

### Figure S11

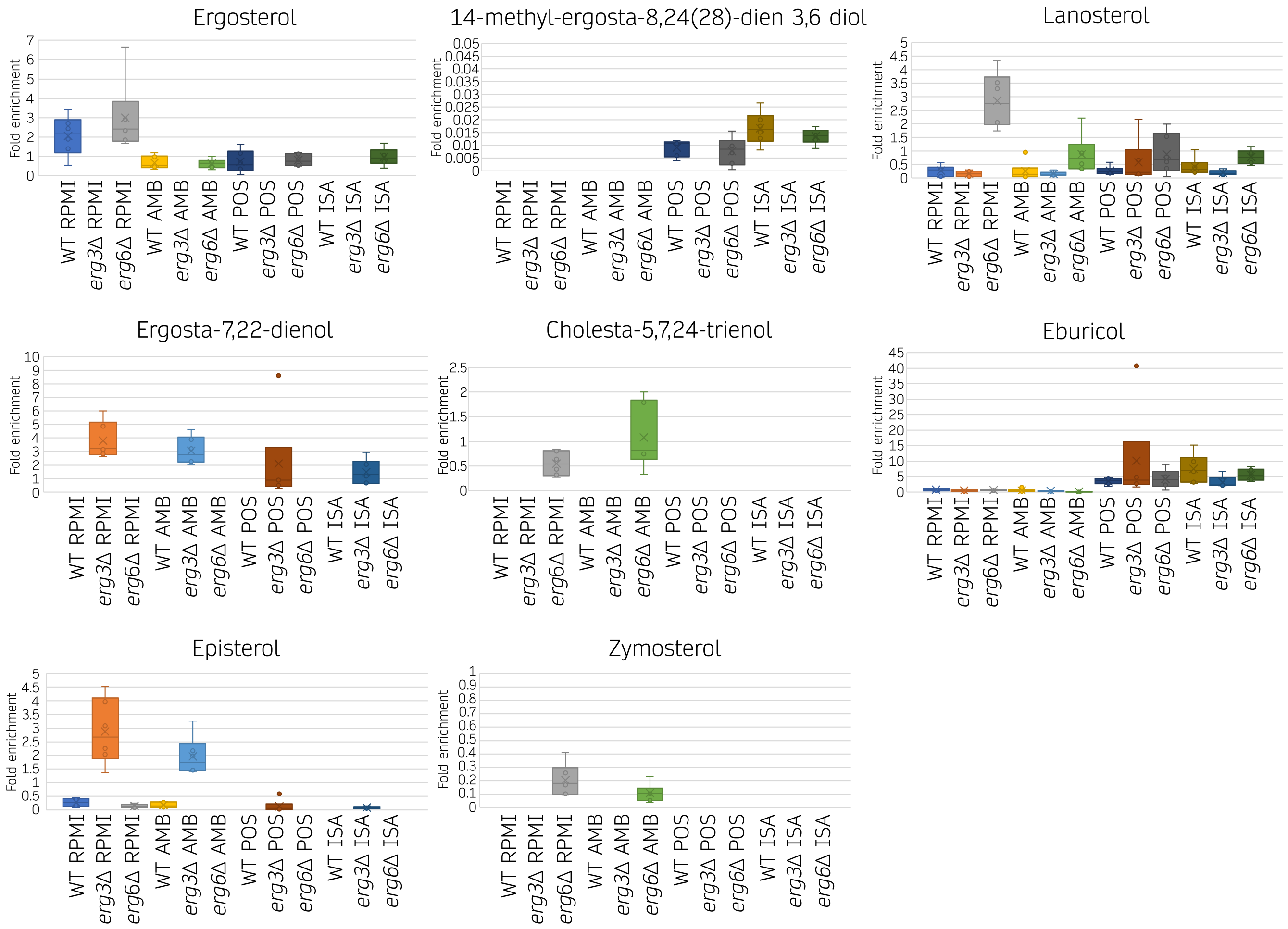


**Figure S11. Normalized sterol abundance in *M. circinelloides erg3*Δ and *erg6a*Δ mutants upon amphotericin B and azole treatments.** Fold enrichment of different sterols content compared to an internal standard and normalized by the dry weight of each sample. Values were determined from 6 biological replicates, 3 from each pair of independently generated mutant for each deletion, to calculate box plots.
